# Functional Predictors of Causative *Cis*-Regulatory Mutations in Mendelian Disease

**DOI:** 10.1101/2020.08.03.232926

**Authors:** Hemant Bengani, Detelina Grozeva, Lambert Moyon, Shipra Bhatia, Susana R Louros, Jilly Hope, Adam Jackson, James G Prendergast, Liusaidh J. Owen, Magali Naville, Jacqueline Rainger, Graeme Grimes, Mihail Halachev, Laura C Murphy, Olivera Spasic-Boskovic, Veronica van Heyningen, Peter Kind, Catherine M Abbott, Emily Osterweil, F Lucy Raymond, Hugues Roest Crollius, David R FitzPatrick

## Abstract

Undiagnosed neurodevelopmental disease is significantly associated with rare variants in *cis*-regulatory elements (CRE) but demonstrating causality is challenging as target gene consequences may differ from a causative variant affecting the coding region. Here, we address this challenge by applying a procedure to discriminate likely diagnostic regulatory variants from those of neutral/low-penetrant effect. We identified six rare CRE variants using targeted and whole genome sequencing in 48 unrelated males with apparent X-linked intellectual disability (XLID) but without detectable coding region variants. These variants segregated appropriately in families and altered conserved bases in predicted CRE targeting known XLID genes. Three were unique and three were rare but too common to be plausibly causative for XLID. We compared the *cis*-regulatory activity of wild-type and mutant alleles in zebrafish embryos using dual-color fluorescent reporters. Two variants showed striking changes: one plausibly causative (*FMR1*^CRE^) and the other likely neutral/low-penetrant (*TENM1*^CRE^). These variants were “knocked-in” to mice and both altered embryonic neural expression of their target gene. Only *Fmr1*^CRE^ mice showed disease-relevant behavioral defects. *FMR1*^CRE^ is plausibly disease-associated resulting in complex misregulation of *Fmr1*/FMRP rather than loss-of-function. This is consistent both with absence of Fragile X syndrome in the probands and the observed electrophysiological anomalies in the *FMR1*^CRE^ mouse brain. Although disruption of *in vivo* patterns of endogenous gene expression in disease-relevant tissues by CRE variants cannot be used as strong evidence for Mendelian disease association, in conjunction with extreme rarity in human populations and with relevant knock-in mouse phenotypes, such variants can become likely pathogenic.

## Introduction

*Cis*-regulatory elements (CRE; encompassing enhancers and repressors) are genomic sequences that control transcriptional activity of one or more genes on the same chromosome via sequence specific interaction with proteins and/or RNA. CRE can be predicted using comparative genomics,^1^ transcriptional characteristics,^2^ patterns of histone modifications and protein association,^3^ patterns of accessible chromatin,^4^ and direct interactions with promoters^5^. Estimates of the number and nature of CRE in the human genome vary with precise definitions but functional ENCODE data has been interpreted as identifying at least 400,000 putative human enhancers^6^. The role of disrupted CRE function in highly penetrant genetic disease was first recognized in association with structural chromosome anomalies which result in loss or gain of regulatory function through deletion or translocation^7, 8, 9, 10^. However the identification of disease associated variants within individual CRE has been complicated by several factors. CRE can function over large genomic intervals and the targeted gene may not be the closest gene. Once a target gene assignment is made the existence of shadow CRE (multiple CRE driving similar expression patterns of the same gene),^11^ can create redundancy and thus tolerance to mutation of individual CRE. A more practical problem is that most CRE exist in the non-coding parts of the human genome where our current understanding of mutation consequence is very incomplete compared to the coding region.

Human developmental disorders provide a powerful system for studying the mechanisms underlying genetic disease in general and regulatory mutations in particular. This diverse group of severe and extreme phenotypes have their onset in embryogenesis or early brain development. Developmental disorders are primarily genetically determined with a high proportion of causative variants arising as *de novo* mutations^12^. Haploinsufficiency is a common mechanism, which affects many different dosage sensitive genes that show complex patterns of expression during development. The genomic intervals encompassing known developmental disorder genes are commonly enriched in highly conserved CRE^13^. There is evidence of enrichment for *de novo* variants in evolutionarily conserved brain active CRE in severe neurodevelopmental disease at a cohort level,^14^ but the confident assignment of variants as causative in affected individuals is difficult^15^.

Here we have attempted to identify all CRE on the X chromosome and then sequence these in 48 individuals with intellectual disability (ID) and a family history that suggests the disease may be X-linked ID (XLID). All affected individuals were recurrently screened negative for likely causative coding mutations on the X chromosome. Using a rational approach to filtering we identified variants in predicted CRE that are associated with known XLID genes and used a range of *in vivo* assays to find features that discriminate likely neutral from likely causative variants.

## Results

### Selection of study cohort

In a previous study we have shown that a significant number of individuals with XLID have no likely disease-associated variants in the coding sequence on the X chromosome,^16^ and subsequent clinical and research analyses. From this group of undiagnosed individuals we identified 48 unrelated males from families with 3 or more affected members with an inheritance pattern strongly suggestive of XLID (Supplementary **Fig. 1**). We reasoned that this cohort should be enriched for regulatory mutations. These families also increase the prior probability that any causative CRE variants would be on the X chromosome thus significantly reducing the genomic search space for interrogation.

### Identification of cis-regulatory enhancers on the X chromosome (chr X)

We have previously identified >100,000 putative CRE covering 4.4% of the human X chromosome and assigned the likely target gene using evolutionary conservation of linkage between CRE and genes located within 1.5 Mb of each other^17^. Approximately a third of CRE could be assigned to a single gene with the remainder having more than one equally plausible target. 389/812 protein coding genes on the X chromosome could be assigned to at least one CRE. Chromatin immunoprecipitation for H3K4me1 in human developing brain showed 10-fold enrichment of these putative CRE. Fluorescent reporter transgenic zebrafish showed >60% of the ~1000 analyzed CRE drive expression in a pattern that overlaps that of endogenous gene activation during development^17^.

### Targeted sequencing and variant filtering of chrx coding regions and enhancers

In the present study we used a targeted sequencing approach to identify variants within all coding regions and CRE on the X chromosome that may be causing XLID in the 48 families using a custom 15.9 Mb oligonucleotide pull-down consisting of 227323 baits. A total of 40,699 variant calls passed basic quality controls in these individuals (**Fig. 1a**). As expected, no clearly disease-associated variants were identified in the coding exons in any of the probands. 628 rare/ultrarare hemizygous variants were identified in high confidence putative CRE of which 31 altered highly conserved bases in enhancers that were predicted to control known XLID genes. 30/31 were confirmed by Sanger sequence analysis in the probands but only 6 of these 30 were shown to segregate appropriately in the XLID families using samples from additional affected, unaffected males and obligate females (**Fig. 1b**). Details of segregation in pedigrees are shown in (Supplementary **Fig. 1 &** Supplementary **Table 4**). 4/48 probands carried one of these six variants and 1/48 carried two.

**Fig. 1:**
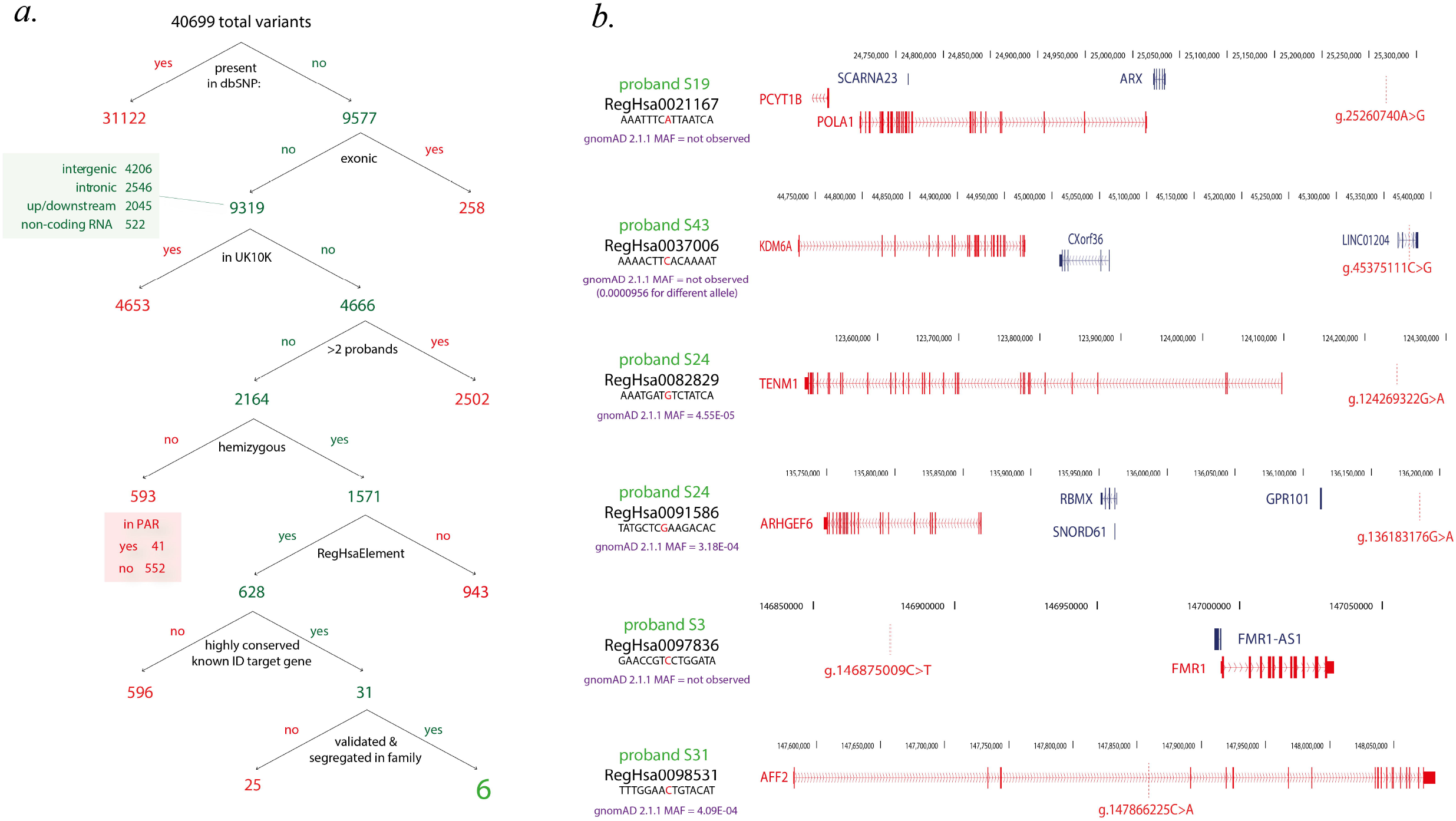
Identification and filtering of XLID-associated regulatory variants and their predicted target genes. **a.** Workflow of the sequencing and curation pipeline leading to the identification of six XLID-associated CRE variants in five probands. **b.** Schematic showing the genomic region of the six genomic variants in the five probands (S19, S24, S3, S43. S31) indicating the location of the XLID-associated CRE variants along with the gnomAD frequencies and their predicted target genes (indicated in red, genomic coordinates form h19/GRCh37 genome build).

### Allele frequencies to assess the plausibility of each rare variant being causative

The position and gnomAD variant allele frequencies (AF) of these six variants are shown in (Supplementary **Table 5**). We used the approach of Whiffin et al,^18^ to calculate the maximum plausible allele frequency for a causal variant in any of the CRE. We chose very conservative parameters: 0.01 for genetic heterogeneity (i.e. 1% of all undiagnosed XLID is caused by a variant in one CRE), 0.2 for allelic heterogeneity (i.e. only 5 different causative variants can exist per CRE) and 0.5 for penetrance (complicated by X-linked inheritance but likely to be ~1 in males and >=0.1 in females). These parameters gave maximum permitted 95% confidence AF = 4e06. Three of the five individuals (S3, S19 and S43) with CRE variants that survived initial filtering do carry variants that may be plausibly disease associated on the basis of the gnomAD AF. The single variant in S31 and both variants in S24 were confirmed to be rare but have gnomAD AF that are too common to be likely disease associated.

### Reporter transgenic analysis of rare filtered variants segregating with the disorder

The reference and alternative base versions of all six rare or ultrarare CRE variants were then tested for CRE function using dual-color fluorescent transgenic assay in zebrafish^19^. In these experiments the mutant CRE drives expression of one fluorescent protein and the wild-type CRE controls a different fluorescent protein in the same fish. Multiple stable lines are created, expression domains are scored and only consistent differences between the reference and alternative alleles are taken as evidence of a functional effect of the mutation. Only two variants in the CRE controlling the XLID genes (in two different probands), *TENM1*^CRE^ and *FMR1*^CRE^, demonstrated a consistent restriction of expression of reporter gene in brain as a result of the variants (**Fig. 2c, d**, 3c, d). The domain of expression of the reporter gene driven by *TENM1*^CRE^ and *FMR1*^CRE^ overlap with the expression domain of the target gene in zebrafish respectively (Fig. 2a, 3a).

**Fig. 2:**
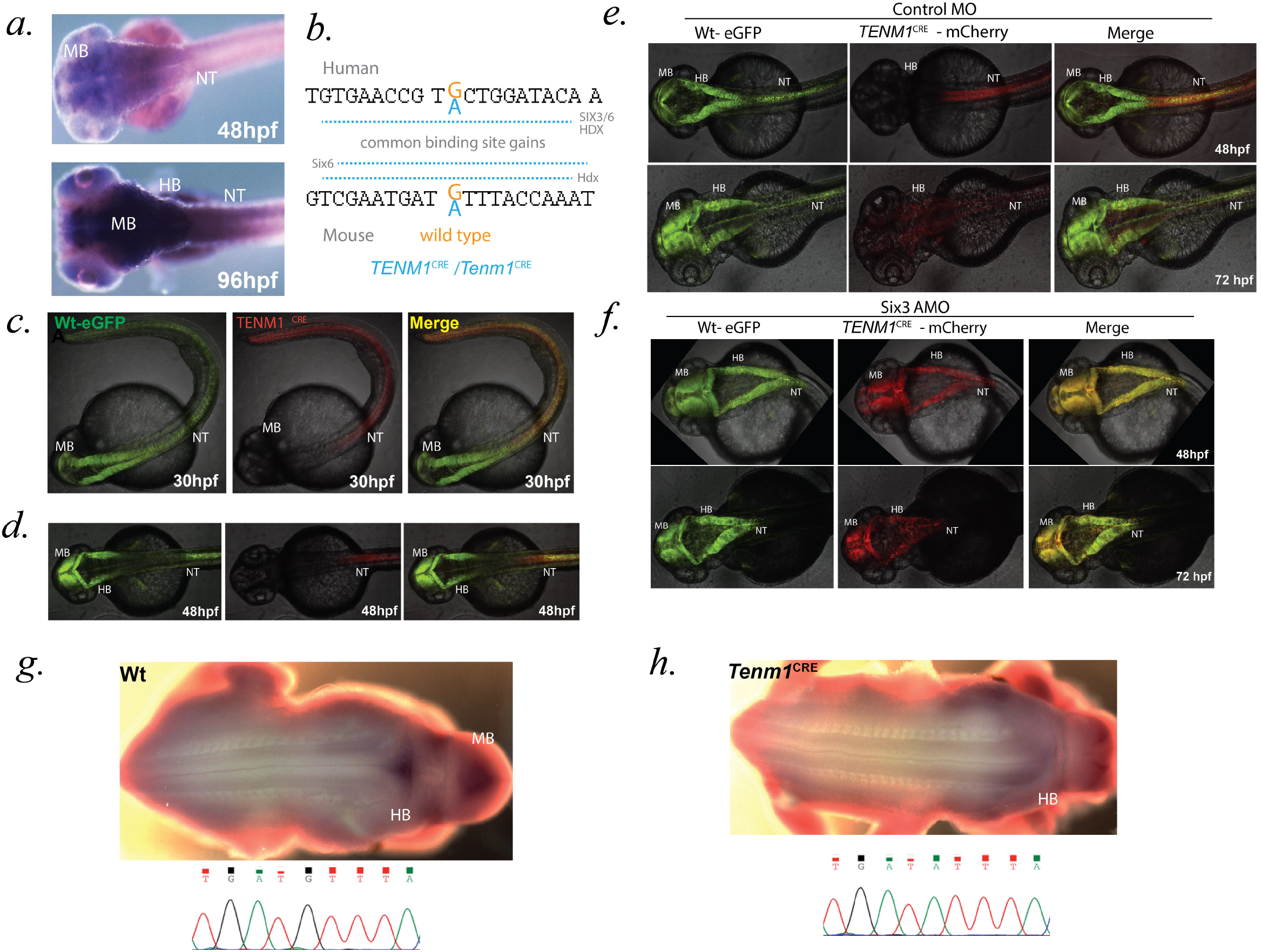
*TENM1* associated regulatory variant alters the expression of *Tenm1* in mouse embryonic development. **a.** *mRNA in situ* hybridization showing expression of *tenm1* in midbrain, hindbrain and neural tube during embryonic development in wild type zebrafish. b. **Human and mouse** (*TENM1*^CRE^ /*Tenm1*^CRE^) **sequences are shown with the variant base marked in blue, resulting in gain of SIX3/SIX6 and HDX binding sites in** *TENM1*^CRE^ **and Six6 and Hdx binding sites in** *Tenm1*^CRE^. **c-d.** Dual color fluorescent transgenic assay in zebrafish with wild-type (Wt) and mutant *TENM1*^CRE^ driving eGFP and mCherry expression respectively. Loss of enhancer activity is observed in midbrain and hindbrain with the mutant *TENM1*^CRE^ allele. **e-f.** *six3* knockdown rescues the effect of the mutant variant on the activity of *TENM1*^CRE^. Control morpholino injected embryos show loss of reporter activity in midbrain and hindbrain by mutant allele, where the mutation creates a Six3 binding site (e). Knock-down of *Six3* rescues the activity of mutant allele in the midbrain and hindbrain (f). **g-h.** Whole-mount *in situ* hybridization for *Tenm1* shows loss of expression of *Tenm1* in the hindbrain and midbrain of *Tenm1*^CRE^ mutant embryos as compared to wild-type embryos. MB: Midbrain; HB: Hindbrain; NT: Neural tube; hpf: hours post fertilization

**Fig. 3:**
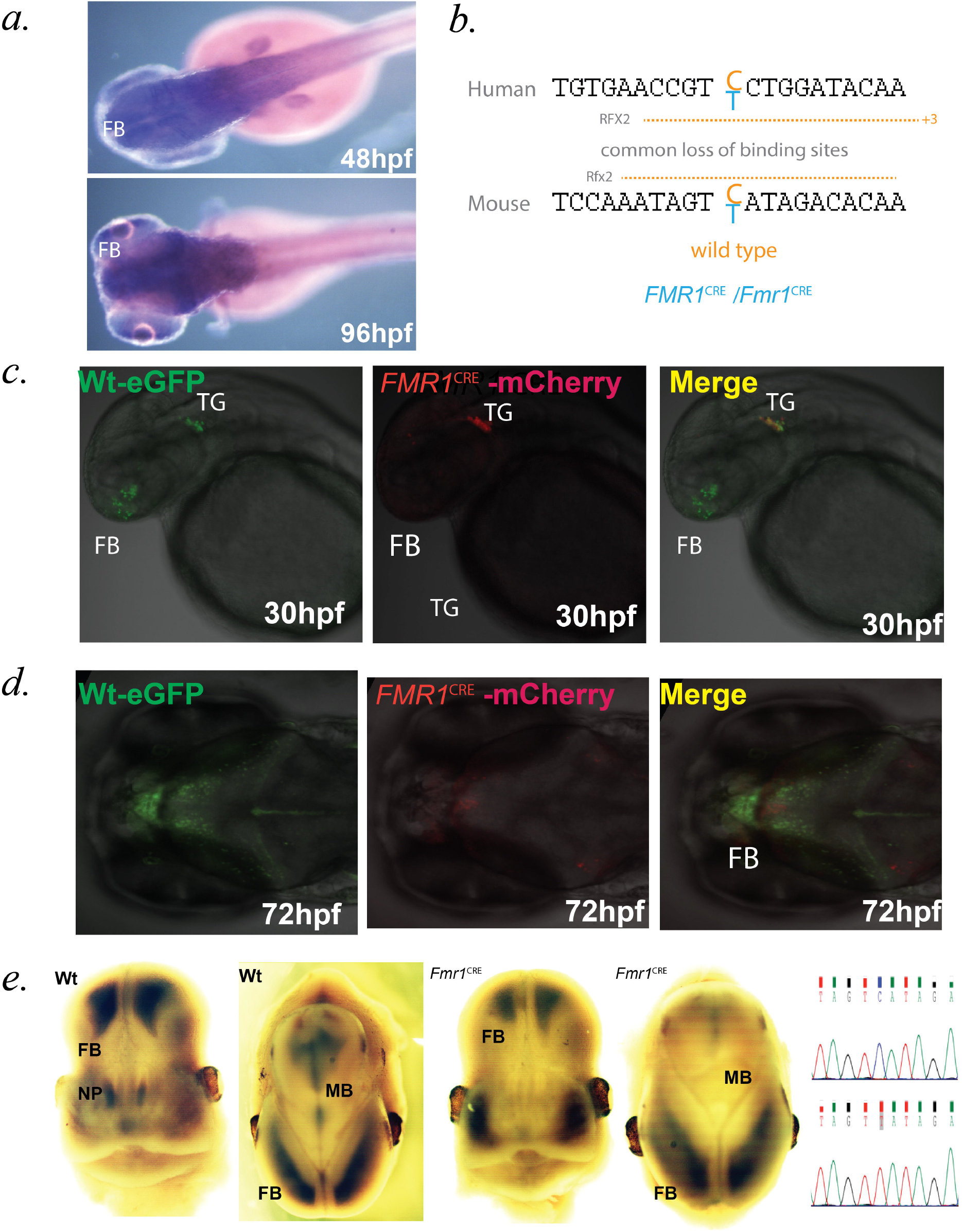
*FMR1* associated regulatory variant alters the expression of *Fmr1* in mouse embryonic development. **a.** *mRNA in situ* hybridization showing expression of *fmr1* in forebrain and midbrain during embryonic development in wild-type zebrafish. b. **Human and mouse** (*FMR1*^CRE^/*Fmr1*^CRE^) **sequences are shown with the variant base marked in blue, resulting in loss of RFX2/Rfx2 binding site in** *FMR1*^CRE^/*Fmr1*^CRE^. **c-d.** Dual color fluorescent transgenic assay in zebrafish with wild-type (Wt) and mutant (Mut) *FMR1*^CRE^ driving eGFP and mCherry expression respectively. Loss of enhancer activity is observed in forebrain with the mutant *FMR1*^CRE^ allele. **e.** Whole-mount *in situ* hybridization for *Fmr1* shows loss of expression of *Fmr1* in the nasal placode and midbrain *Fmr1*^CRE^ mutant embryos as compared to wild-type embryos. FB: Forebrain; MB: Midbrain; TG: Trigeminal ganglia; NP: Nasal placode; hpf: hours post fertilization

### TENM1^CRE^ creates a de novo and functionally repressive binding site for *six3*

We had noted that the *TENM1*^CRE^ variant created a predicted binding site for the homeodomain-containing DNA binding proteins SIX3 or SIX6 in the human element (Fig. 2b). We chose SIX3 for further study as it is essential for early brain development and with pathway-specific activator and repressor activity^20^. To determine if SIX3-mediated repression may be responsible for the altered enhancer activity in the variant *TENM1*^CRE^ we titrated morpholinos against zebrafish *six3* into the embryos from a *TENM1*^CRE^ dual color fluorescent transgenic line to the point where there was no morphological anomaly. This resulted in an alteration in the expression of the *TENM1*^CRE^ transgene to match the wildtype in the morphant embryos (**Fig. 2e, f**), supporting acquisition of SIX3 repression as the mechanism for the transcriptional effect in zebrafish embryos.

### *Fmr1* expression and behavioral phenotypes in *Fmr1*^CRE^ and *Tenm1*^CRE^ mouse models

CRISPR/Cas9 genome editing was used to knock the exact mutation into mouse embryos via homologous recombination for both variants that showed a consistent functional consequence in the zebrafish lines: *FMR1*^CRE^ and *TENM1*^CRE^. We established multiple independent mouse lines for each CRE variant on a C57BL/6 background (Fig. 2b, 3b). All lines resulted in hemizygous mutant animals, at the expected ratio that were healthy and fertile with no obvious morphological abnormalities. We first looked for alteration in the expression of the predicted target gene during development. *Tenm1*^CRE^ resulted in loss of expression in the hindbrain of the mutant embryos using whole-mount *in situ* hybridization (WISH) for the endogenous gene (**Fig. 2g, 2h**). *Fmr1*^CRE^ caused a significant reduction in the developmental expression of *Fmr1* in the olfactory placodes and the forebrain (**Fig. 3e**). Both variants have an effect on endogenous gene expression i.e. neither is behaving as a shadow CRE.

To look for functional phenotypic effects segregating with either CRE variant we first tested olfaction. This sense was selected for two reasons. First, the complete loss of *Fmr1* expression in the olfactory placode in *Fmr1*^CRE^ embryos. Secondly, *TENM1/Tenm1* mutations have recently been identified in humans and mice associated with congenital generalized anosmia^21^. Using a buried chocolate button test hemizygous *Fmr1*^CRE^ mice showed a significant increase in time to discovery compared to wild-type male littermates (**Fig. 4g**). *Tenm1*^CRE^ hemizygotes had olfactory function similar to wild-type male littermates (**Fig. 4h**).

**Fig. 4:**
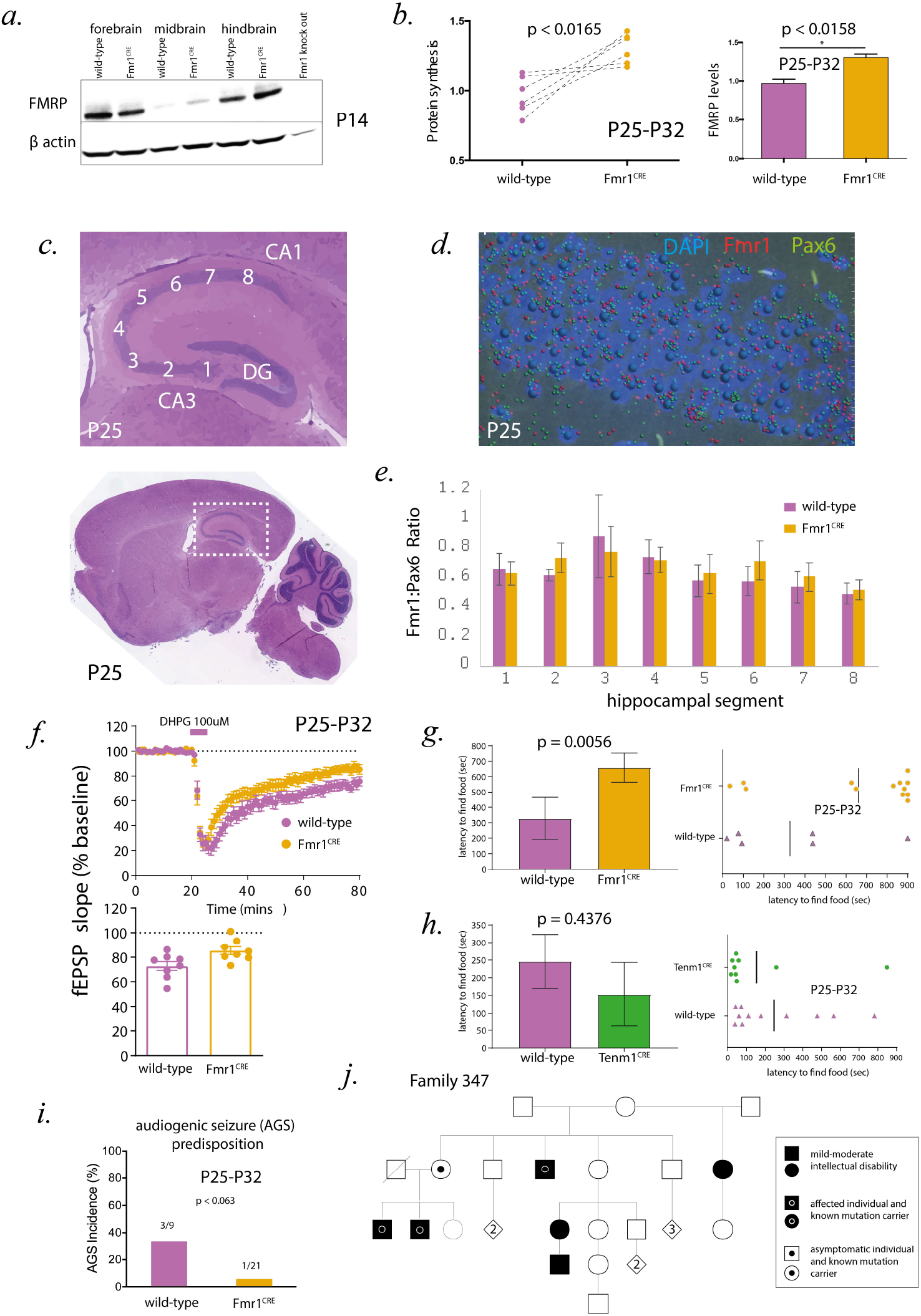
Functional phenotypic effects observed in mice bearing *FMR1* associated regulatory variant. a) Levels of FMRP protein observed in the forebrain, midbrain and hindbrain of *Fmr1*^CRE^ knock-in mutant mice as compared to wild-type litter mates at P-14. b) Significant increase in bulk protein synthesis levels in slices from dorsal hippocampus of *Fmr1*^CRE^ knock-in mutant male mice as compared to wild-type male littermates. c) H&E stained brain sagittal section with marked hippocampus regions is shown as reference image on which RNAscope analysis was done *Fmr1*^CRE^ mutant male mice compared to wild-type male littermates. Images of RNAscope processed CA3-CA1 components of the hippocampus brain section were taken as shown by numbers (1-8) starting from dentate gyrus (DG). d) Reference image of RNAscope processed section with *Fmr1* transcript (Red), *Pax6* transcript (Green) and nucleus (Blue/DAPI). Each transcripts are represented by spots. e) Graphical representation of *Fmr1* transcripts normalised to *Pax6* transcripts (used as control) is shown across the whole CA3-CA1 components of the hippocampus brain section between *Fmr1*^CRE^ mutant male(purple) mice compared to wild-type littermates(orange). No significant difference was overserved in the *Fmr1* transcripts level across the whole region used for analysis. f). mGluR-dependent LTD is significantly decreased in CA3-CA1 components of the hippocampus of *Fmr1*^CRE^ mutant mice as compare to wild type litter mate **(g-h) Olfactory function.** The mice hemizygous for the variant in *Fmr1*^CRE^ showed a significant increase in time to discovery compared to wildtype male controls in a buried food test. No significant difference in the levels of latency to find food was observed in mice hemizygous for the variant in *Tenm1*^CRE^ compared to wild type litter mates. i) **Audiogenic seizures.** No significant difference was observed in audiogenic seizure incidence in the hemizygous mice with the variant *Fmr1*^CRE^ compared to wild-type littermates. j) Pedigree of Family 347 of which individual S3 is a member showing segregation of the mutation affecting *FMR1* expression.

### Abnormal hippocampal protein synthesis and electrophysiology in *Fmr1*^CRE^ mice

*FMR1*/*Fmr1* encodes 516-622 amino acid RNA-binding and polyribosome associated protein isoforms (FMRP) that are essential for the normal development and function of neurons in the brain. Loss of FMRP function is responsible for Fragile X syndrome, the most common form of XLID. A well-characterised biochemical effect of loss of FMRP in the brain of the mouse model of Fragile X syndrome is the mGluR5- or ERK1/2-dependent elevation of basal protein synthesis^22^. We found a significant increase in bulk protein translation levels in tissue slices of dorsal hippocampus of *Fmr1*^CRE^ mutant male mice compared to wild-type male littermates (**Fig. 4b**). This would be consistent with the reduction of *Fmr1* expression we observed in mutant embryos (**Fig. 3e**). Importantly we did not find significant difference in the expression of *Fmr1 transcript* in *Fmr1*^CRE^ mutant male mice compared to wild-type male littermates at developmental stage P-7 (by qPCR), P-14 (by qPCR) (**Supplementary Fig. 2**) or P-25 (by RNAScope) (**Fig. 4c,d,e**) and (by RNA Sequencing) (**Supplementary Fig. 3**).

Given the gene expression results, it was surprising to find an increase in FMRP protein abundance in the hippocampus of *Fmr1*^CRE^ mutant male mice as compared to wild-type litter mates using western Blotting (**Fig. 4a, b; Supplementary Fig. 4**). The increase in FMRP protein levels is consistent with our finding that mGluR-dependent long-term depression (LTD) in the CA3-CA1 regions of the mouse hippocampus is significantly decreased in *Fmr1*^CRE^ hemizygous mutant male mice as compared to wild-type litter mates (**Fig. 4f**) as this is exaggerated in *Fmr1*-null animals^23^. *Fmr1*^CRE^ hemizygous mutant mice were also found to have no increase in audiogenic seizure predisposition, a phenotype that typifies *Fmr1* null mice (**Fig. 4i**). These latter three key features strongly suggest that *Fmr1*^CRE^ does not represent a simple loss of FMRP function.

### Clinical re-evaluation and whole genome sequencing of individuals carrying *FMRI*^CRE^ CRE

Re-evaluation of affected individuals within the family in which *FMR1*^CRE^ is segregating (Fig. 4j) revealed no features suggestive of a Fragile X (FRAX) syndrome diagnosis (OMIM #300624]; FMRP deficiency) other than macrocephaly and intellectual disability. Importantly none of the individuals carrying *FMR1*^CRE^ showed clinical features of FRAX Tremor and Ataxia Syndrome (FRAXTAS [OMIM #300623]; FMRP over-production)^24^. Whole genome sequencing of individual S3 (*FMR1*^CRE^ proband) did not identify any other plausible cause of his intellectual disability.

## Discussion

The motivation for initiating this study was the difficulty in assigning pathogenic or likely pathogenic status to a *de novo* or segregating variant in a regulatory sequence. Currently almost all such ultra-rare variants would be considered of uncertain significance using current best practice guidelines^25, 26^. However it has been shown that using “well-established” functional assays demonstrating a variant has abnormal gene function (coded as assigning PS3 in the guidelines) has the potential to change many variants of uncertain significance (VUS) to likely pathogenic status^27^. The question then becomes: how should we use data from functional assays in clinical interpretation of regulatory variants. Given the rapid switch from targeted whole exome sequencing to whole genome sequencing it is likely that there will be an increasing need to develop a rational approach to the interpretation of ultra-rare regulatory variants.

Here we attempted to perform an integrated clinical, genetic, developmental, behavioural and neurophysiological approach to the analyses of CRE variants identified in a cohort of affected individuals with a Mendelian phenotype that should be enriched for causative cis-regulatory mutations. XLID accounts for ~16% of ID in males^28^. Mutations in the coding region of at least 81 different genes,^16, 29^ have been identified as causing XLID. Given the significant contribution of XLID to ID and the observed regulatory variant enrichment in a large cohort of individuals with neurodevelopmental disorders,^14^ we reasoned that we could increase the prior probability of identifying likely causative mutations by restricting the genomic search space to the X chromosome and by limiting our investigations to variants in enhancers that targeted known XLID genes. This strategy was implemented as most known disease-associated regulatory mutations were identified because they partially,^30^ or fully,^31^ phenocopy loss-of-function mutations in the target gene. If true for our cohort, then matching the pattern of clinical features of individuals carrying a specific regulatory mutation to those of the syndrome associated with intragenic mutations would have diagnostic value.

One important feature of our study was that we could compare the functional impact of variants which were plausibly responsible for a significant Mendelian disorder and those that were too common to be disease associated using established statistical approach^18^. The *in vivo* analysis of the *TENM1* enhancer was particularly interesting in this regard. Both the transgenic zebrafish embryos and knock-in mouse lines showed clear evidence of abnormality in developmental gene expression. Indeed, we were able to identify the transcription factor mediating the repressive effect of the rare CRE variant in zebrafish. This could be taken as strong evidence of disease association if the significance of the allele frequency was not appreciated. It seems likely that simpler and more commonly used *in vitro* assays of enhancer activity,^32^ will be more prone to “over-reporting”. It is also clear that most unique variants will also be neutral or of low-penetrant effect so the allele frequency on its own is not sufficient for discrimination in clinical analysis.

The unique CRE variant *FMR1*^CRE^ is the most plausible disease associated allele of those identified in this study. This variant produced abnormal embryonic expression of endogenous *Fmr1* in the mouse model (**Fig. 3e**). The reporter transgenic analysis in zebrafish also showed an expression pattern that would be consistent with tissue specific loss of function during early brain development. In contrast, we were unable to show evidence of significant transcriptional misregulation in the *Fmr1*^CRE^ post-natal brain using qRT-PCR (Supplementary **Fig. 2**), RNA-Seq (Supplementary **Fig. 3**) or RNAscope *in situ* hybridization (**Fig. 4d, e**). This was particularly interesting in the context of the other analyses that we performed in P25 *Fmr1*^CRE^ mice. First the pattern of bulk protein synthesis was similar to that seen in *Fmr1* KO mice. However, the direction of electrophysiological change of LTD was opposite from *Fmr1* KO mice and increased levels of FMRP protein were observed **(Fig. 4a, b**; Supplementary **Fig. 4**). These apparently paradoxical results may be the consequence of developmental mis-programming of the cells in the hippocampus. In the future we intend to employ single cell transcriptomics and ATAC-Seq to test this hypothesis.

We chose the buried food behavioural assay to assess olfaction because this would be plausibly disrupted in both *Tenm1*^CRE^^21^, and *Fmr1*^CRE^^33, 34^. Like all mouse behavioural assays this is very likely to be influenced by other neurodevelopmental and environmental factors. That said, the fact that it was normal in *Tenm1*^CRE^ and impaired in *Fmr1*^CRE^ is probably significant given the striking loss of *Fmr1* expression in the olfactory placodes in *Fmr1*^CRE^ embryonic mice. Surprisingly, given the human phenotype, it has been difficult to identify any behavioural assay that provides consistent evidence of cognitive impairment in *Fmr1* KO mice^35^.

For the reasons outlined in the two paragraphs above we do not consider it to be surprising that the proband S3 and his affected male relatives carrying *FMR1*^CRE^ do not show a clinical pattern typical of either Fragile X syndrome [OMIM 300624] or FRAXTAS [OMIM 300623]. The family presented with a non-specific intellectual disability associated with mild macrocephaly. We consider it likely that many disease associated CRE variants will result in clinical features that significantly differ from those seen associated with intragenic mutations of target gene. If this heterogeneity in clinical features is indeed true, the clinical genetics field will have relatively limited ability to predict the phenotypes associated with regulatory mutations even when the clinical impact of intragenic mutations of target gene are well characterised.

We can conclude that although it remains challenging to recognise highly penetrant CRE causative variants, the population allele frequency estimates from gnomAD 2.1.1 has power to identify those of neutral or low penetrant effect. These data allowed us to class *TENM1*^CRE^ as implausible as an XLID causative variant despite it being in an evolutionarily conserved, non-redundant CRE with a strong repressive effect on *Tenm1* expression during mouse hindbrain development. The discriminative power of the extreme rarity of individual alleles may prove particularly useful for the identification of causative variants in CRE which are poorly conserved across species but under high levels of selective constraint within human populations^36^. For the time being we suggest that causative variants are restricted to those with strong human genetic evidence, supported by modelling of the precise variant *in vivo* and resulting in both transcriptional misregulation and phenotypic effect.

## Materials and methods

### Cohort Selection

Genomic DNA samples from 48 individuals (probands) with moderate-to-severe intellectual disability (ID) were used in this study. The appropriate research ethical approval was obtained (IRAS 03/0/014), and parents or guardians provided informed written consent. Each individual is assumed to have X-linked recessive form of ID on the basis of positive family history: three or more cases of ID in males only, predominant sparing of carrier females and no evidence of male-to-male transmission of the disease (Supplementary Fig. 1). A clinical geneticist had assessed the individuals and the cause of the ID was unknown. The severity of the disease was categorized using DSM–IV or ICD-10 classifications (profound mental retardation was classified as severe). The patients had previously been tested negative by routine diagnostic approaches (i.e., CGH microarray analysis at 500 kb resolution, fragile X [MIM #300624], methylation status of Prader Willi [MIM #176270]/Angelman syndrome [MIM #105830]). In addition, coding variants on the X chromosome likely to lead to disease have not been found within a previous study^16^.

### Targeted Capture Design and Sequencing

A comprehensive list of coordinates of all the exonic and conserved regulatory elements from human X chromosome was used to design a customized capture library from Roche, NimbleGen (Supplementary Table 1). Library preparation, pre and post capture multiplexing were performed using the SeqCap EZ Choice XL kit (Roche NimbleGen) and TruSeq index barcodes (Illumina) were used according to the manufacturer’s instructions. 4 different DNA samples were pooled for pre capture multiplexing and 4 post captured libraries were combined and paired-end sequenced performed on a single lane of a HiSeq-2000 instrument (Illumina). In total 16 different DNA samples were sequenced in a single lane of a HiSeq-2000 and 4 lanes were used to sequence all the 48 DNA samples.

### Read Mapping, Variant Analysis and Enhancer Selection

Following quality control with FastQC, reads were mapped to the GRCh37 version of the human reference genome using BWA^37^. Variants were called using GATK,^38^ according to its recommended best practice pipeline. 40,699 variants remained after filtering out variants that failed GATK’s variant quality score recalibration. These variants were subsequently compared to dbSNP v137 to filter out common variants. Any variant with one of the following handles in dbSNP (1000GENOMES, CSHL-HAPMAP, EGP_SNPS, NHLBI-ESP, PGA-UW-FHCRC) were excluded where the variant’s reported minor allele frequency was greater than 0.01 and the minor allele was reported to be observed in at least two samples. The remaining 9,577 X chromosome variants were then annotated with SnpEff,^39^ to determine their predicted effects on genes. gnomAD 2.1 allele frequencies were documented for the surviving variants. To determine the best candidates for experimental validations, the variants were ranked based on extreme evolutionary conservation. Using Multiple Sequence Alignments from 45 vertebrate species against the Human genome (UCSC genome browser), mutations were retained if the reference human allele was conserved in at least 90% of the species, and then sorted by decreasing conservation depth. Top variants were then manually evaluated using biochemical signals from the ENCODE project (H3K4me1, H3K4me3, H3K27ac, DNase1 sensitivity), and based on the association to target genes known to be responsible for XLID or functionally related to brain development, leading to a final selection of 31 candidate variants (Supplementary **Table 4**). Target genes for each of the CRE harbouring the variants were assigned as described previously^17^. Motif search on CRE element was performed on a 40bp window around the mutated base for both human and mouse sequences using the FIMO software from the MEME suite^40^. The motif databases used for the search were Jaspar Core 2018 for vertebrates and Uniprobe mouse motifs as downloaded from the MEME website. Motifs with a p-value of 0.001 or lower that were present uniquely in either the WT or the mutant sequences are reported.

### Transgenic Zebrafish, In Situ Hybridization (ISH) and Morphant Generation

All mouse and zebrafish experiments were approved by The University of Edinburgh ethical committee and performed under UK Home Office license number PIL 60/12763, 70/25905, I655D57B6, PA3527EC3 and 1724D1B2C; PPL 60/4418, 60/4424, IFC719EAD and 60/4290. The wild type and mutant versions of the *FMR1*^CRE^ and *TENM1*^CRE^ were analyzed for their regulatory activities in dual color enhancer-reporter transgenic assays in zebrafish embryos^19^. The sequences of the primers used in generating the constructs utilized in the assay are listed in (Supplementary **Table 2).** A summary of the number of independent lines analyzed for each enhancer and their expression sites is included in (Supplementary **Table 3**). The transgenic F1 embryos were processed for imaging as described^19^. The images were taken on a Nikon A1R confocal microscope and processed using A1R analysis software. A zebrafish *six3* antisense morpholino oligonucleotide (Six3AMO) was obtained from Gene Tools, LLC, with the following sequence: 5’ GCTCTAAAGGAGACCTGAAAACCAT 3’. This morpholino has sequence complementary to the highly conserved sequences around the translation initiation codon of both *six3a* and *six3b*, and hence inhibits the function of both zebrafish *six3* genes^41^. As control we used the Gene Tools LLC standard negative control morpholino: 5’ CCTCTTACCTCAGTTACAATTTATA 3’. The morpholinos were injected into 1 to 2-cell stage of at least 100 embryos to deliver an approximate amount of 2.5 ng per embryo. RNA in situ hybridization on fish embryos was performed as previously^42^. The sequences of primers used for synthesis of specific probes are listed in (Supplementary **Table 2**).

### Generation of Transgenic Mice and Embryo ISH

CRISPR/Cas9 gene targeting technology was used to generate mouse lines with orthologous mutations; *Fmr1*^CRE^ and *Tenm1*^CRE^. Double stranded DNA oligomer that provides a template for the guide RNA sequence was cloned into px461. The details of guide RNA and repair template sequence are in **Supplementary note.** The full gRNA template sequence is amplified from the resulting px461 clone using universal reverse primer and T7 tagged forward primers. The guide RNA PCR template is used for *in vitro* RNA synthesis using T7 RNA polymerase (Neb), and purified using RNeasy mini kit (Qiagen) purification columns. The zygotic injection mix contains Cas9 mRNA (Tebu Bioscience @ 50ng/μl), guide RNA (25ng/μl) and repair template single stranded DNA (IDT 150ng/μl). Injected embryos were transferred into the oviducts of pseudopregnant females to litter down. Genotyping of the resulting mice was performed by Sanger sequencing using tail tip DNAs. F0 mice with desired variant were crossed with C57BL/6 to generate a stable mice line. *In situ* hybridization on mouse embryos was performed with DIG-labelled gene-specific antisense probes as previously described^43^. The sequences of primers used for synthesis of specific probes are listed in (Supplementary **Table 2**).

### Behavioral Testing of Fmr1 and Tenm1 Mouse Lines

Male WT (wild type) and mutant (*Fmr1*^CRE^ and *Tenm1*^CRE^ knock-in) littermates were used for the test at P25-32 using the buried food test assay. For three consecutive days before the test, ¼ Cadbury’s chocolate button was placed in the home cage for 15 minutes to habituate the mice to the food reward. 12 hours before the test, all food was removed from the home cage to motivate the mouse to find the food reward during the test. After 12 hours, the mouse was placed in a clean cage with fresh bedding in which ¼ chocolate button had been buried 1cm beneath the bedding. The time taken for the mouse to find the buried food was scored and the test was stopped if the mouse did not find the food after 15 minutes. The bedding was replaced and the cage cleaned with 1% Conficlean between mice. All mice were scored blind to the genotype. Unpaired t-tests were used to determine statistical significance.

### Seizure Propensity Testing of Fmr1^CRE^

Male WT and *Fmr1*^CRE^ knock-in mutant littermates (P25-32) were tested for audiogenic seizures as described previously^44^. Briefly, animals were transferred to a transparent plastic test chamber and, after 1 minute of habituation, exposed to a 2 min sampling of a modified personal alarm held at > 130dB. Seizures were scored for incidence (seizure/no seizure) and severity, with an increasing scale of 1=wild running, 2=clonic seizure, and 3=tonic seizure. All mice were tested and scored blind to genotype. Statistical significance for incidence was determined using two-tailed Fisher’s exact test.

### Basal Protein Synthesis and FMRP Western Blotting

Protein synthesis levels were measured following the protocol outlined by Osterweil^22^. The detailed protocol is described in the **Supplementary note**.

For western blots, forebrain, midbrain and hindbrain from (P-14) and hippocampal slice from P-25 male WT and *Fmr1*^CRE^ knock-in mutant littermates were dissected and homogenized in lysis buffer (20 mM HEPES pH 7.4, 0.5% Triton X-100, 150 mM NaCl, 10% glycerol, 5 mM EDTA with protease inhibitor cocktail (Roche), incubated at 4 °C for 30 min followed by centrifugation at 14000 rpm for 30 min to collect the supernatant. These samples were directly used for SDS-PAGE and transferred onto nitrocellulose membranes for immunoblot analysis with FMR1 (MAB2160, Milipore) and Actin antibodies (13E5, CST).

### Hippocampal Slice Electrophysiology

Electrophysiology experiments were performed as outlined by Stephanes^45^. The detailed protocol is described in the **Supplementary note**.

### RNAscope Assay and Imaging

In situ RNA hybridization was performed using the RNAscope assay (Advanced Cell Diagnostics, ACD, Hayward, CA, USA) according to the manufacturer’s recommendations. The detailed protocol is described in the Supplementary note. The images of sections were processed using the multimodal Imaging Platform Dragonfly (Andor Technologies, Belfast, UK) using air 40x Plan Fluor 0.75 DIC N2. Data were collected in Spinning Disk 25 μm pinhole mode on the high sensitivity iXon888 EMCCD camera. According to Advanced Cell Diagnostics, each mRNA molecule hybridized to a probe appears as separate small puncta. Data visualization and spot counting was done using IMARIS 8.4 (Bitplane).

## Acknowledgements

DRF and VvH were supported by MRC University Unit grant to the MRC Human Genetics Unit at the University of Edinburgh. HB & MN and project costs were supported and funded by the 7th framework programme of the European Union [NeuroXsys Project HEALTH-F4-2009-223262]. HB was subsequently funded by a grant from NewLife (Grant Ref: 14-15/07). National Institute of Health Research Bioresource for Rare Diseases (grant number RG65966) for whole genome sequence data from 12,596 X chromosome alleles as controls. JH is funded by a BBSRC studentship. FLR and DG are funded by NIHR Cambridge Biomedical Research Centre grant. HRC received support from the French Government from programs implemented by ANR with the references ANR–10–LABX–54 MEMOLIFE and ANR–10–IDEX–0001–02 PSL* Research University. DRF,PK and EO receive grant funding from the Simons Initiative for the Developing Brain.

## Author Contributions

DRF, HRC, FLR and VvH conceived the project and with CA and EO designed the experimental approaches. HB, DG, OSB, SB, SRdL, JH and SB performed the experiments and interpreted the results. LM, MN and HRC performed the computational genomic analysis. DRF, HRC and FLR wrote the manuscript with contributions from HB, SB, EO and VvH.

