## Supplementary Data for "Functional Predictors of Causative *Cis*-Regulatory Mutations in Mendelian Disease"

| **Primer details for making transgenic zebrafish** | |
| --- | --- |
| ***TENM1* CRE** **F** | AAGAGAAAAGTGGATGGGAGAGA |
| ***TENM1*CRE** **R** | TCATCTTTATTGAGAAGAATATGGGAC |
| ***FMR1* CRE** **F** | ATTAAGATTCAAGTTCGTTATAGCACT |
| ***FMR1*CRE** **R** | GAAAGTGATATGACAATGGCAACT |
| **Genotyping primers for transgenic mice** | |
| ***TENM1* CRE** **F** | GCTCTCAAATATTGTGTGACAAAG |
| ***TENM1*CRE** **R** | ACCATCACAGGTAGACCTCCTA |
| ***FMR1* CRE** **F** | GAAGAAGCACTCTCCTAAAATGG |
| ***FMR1*CRE** **R** | GCCTTCTCCCTGGTTTATCC |
| **qRT-PCR primers** | |
| ***Fmr1* 6 F (exon14-15)** | CGCGGTCCTGGATATACTTC |
| ***Fmr1* 6 R (exon14-15)** | CTCTGCGCAGGAAGCTCT |
| ***Fmr1* 77 F (exon5-6)** | GATTCCATTCCATGATGTGAGA |
| ***Fmr1* 77 R (exon5-6)** | GCTCTTTTTCATTTGCTCTGG |
| ***Gapdh* F** | GGGTTCCTATAAATACGGACTGC |
| ***Gapdh* R** | CCATTTTGTCTACGGGACGA |
| **Insitu probe primer sequence for zebrafish and mouse** | |
| ***zfodz1* F** | TTTAGAGTGGCCCACAGACC |
| ***zfodz1* R** | GGAGACTTCAGCTTGGCATC |
| ***zfFmr1* probe** | Gift from Steve Wilson Lab |
| ***mFmr1* F** | GCAGCTTGCCTCAAGATTTC |
| ***mFmr1* R** | AGCCTTGGGTTCAGGTTTCT |
| ***mOdz1* F** | CATCACCTGACCATGCACTC |
| ***mOdz1* R** | TGGGTGGTGGTGAGTAGACA |

**Supplementary Table 2: Detail list of primers used in the study.**

| Transgene * | Reporter used in transgenic assay ^#^ | Total number of stable transgenic lines analysed | Sites of reporter expression  driven by the element | Tissue-specific activity of the CRE observed in 100% of transgenic lines analysed |
| --- | --- | --- | --- | --- |
| *TENM1*^CRE^-WT | eGFP | 4 | **Neural tube**  **(4/4; 100%)**  **Hindbrain**  **(4/4; 100%)**  **Midbrain**  **(4/4; 100%)**  Pectoral fin  (2/4; 50%)  Heart (1/4; 25%) | Neural tube, hindbrain, midbrain |
| *TENM1*^CRE^ -Mut | mCherry | 4 | **Neural tube (4/4; 100%)**  Hindbrain  (0/4; 0%)  Midbrain  (0/4; 0%)  Otic vesicle (1/4; 25%)  Olfactory placode (1/4; 25%) | Neural tube |
| *FMR1*^CRE^ -WT | eGFP | 4 | **Forebrain**  **(4/4; 100%)**  **Trigeminal ganglia**  **(4/4; 100%)**  **Lateral spinal cord neurons**  **(4/4; 100%)**  Heart (1/4; 25%) | Forebrain, trigeminal ganglia and lateral spinal cord neurons |
| *FMR1*^CRE^-Mut | mCherry | 4 | Forebrain  (0/4; 0%)  **Trigeminal ganglia**  **(4/4; 100%)**  **Lateral spinal cord neurons**  **(4/4; 100%)**  Neural tube (1/4; 25%) | Trigeminal ganglia and lateral spinal cord neurons |

**Supplementary Table 3:** **Transgenic lines generated in zebrafish**. Details of Wild type and variant CRE (*TENM1* and *FMR1)* driven transgene expression for sites in F1 embryos obtained from multiple independent stable transgenic F0 lines **(* Species of origin: *Homo sapiens* # Model organism for reporter transgenic assay: *Danio rerio)***

| Proband | Variant | Target Gene | Alleles | Total Alleles | Hemizygotes | Allele Frequency* |
| --- | --- | --- | --- | --- | --- | --- |
| S3 | X-146875009-C-T | *FMR1* | 0 | NA | 0 | 0 |
| S19 | X-25260740-A-G | *POLA1 PCYT1B* | 0 | 21967 | 0 | 0 |
| S24 | X-124269322-G-A | *TENM1* | 1 | 21962 | 0 | 0.00004553 |
| S24 | X-136183176-G-A | *ARHGEF6* | 7 | 21979 | 1 | 0.0003185 |
| S31 | X-147866225-C-A | *AFF2* | 9 | 22020 | 3 | 0.0004319 |
| S43 | X-45375111-C-G | *KDM6A* | 0 | NA | 0 | 0 |

***Popmax Filtering AF (95% confidence)**

**Supplementary Table 5: gnomAD 2.1.1 Frequencies of six CRE variants surviving initial filters.**

##### Metabolic labeling and Basal Protein Synthesis. Juvenile (P25-P32) male littermate WT and *FMR1^CRE^* knock-in mutant mice were anesthetized with isofluorane, and the hippocampus dissected into ice-cold artificial cerebral spinal fluid (ACSF) (in mM: NaCl: 124, KCl: 3, NaH2PO4: 1.25, NaHCO3: 26, dextrose: 10, MgCl2: 1, CaCl2: 2, saturated with 95% O2 and 5% CO2). Slices (500 µm thick) were prepared using a Stoelting Tissue Slicer, and within 5 min transferred into 32.5°C ACSF (saturated with 95% O2 and 5% CO2) for 3.5–4 h to allow for recovery of protein synthesis as described^1^.25 µM ActD was then added for 30 min to inhibit transcription. Slices were incubated in 10 µCi/ml 35S-Met/Cys (express protein labelling mix, Perkin Elmer) for another 30 min to measure protein synthesis. Slices were homogenized in ice-cold homogenization buffer (10 mM HEPES pH 7.4, 2 mM EDTA, 2 mM EGTA, 1% Triton X-100, protease inhibitors tablets (Roche), and phosphatase inhibitors (cocktails 1+2,Sigma Aldrich), and precipitated using trichloroacetic acid (TCA; 10% final) for 10 min on ice and pelleted by spinning at 21,000×g for 10 min. The pellet was washed with ice-cold ddH2O and resuspended in 1 N NaOH then adjusted to a neutral pH with HCl. Triplicate aliquots were added to the scintillation cocktail (Optiphase, Perkin Elmer) and read with a scintillation counter. Averaged triplicate counts per minute (CPM) per µg protein were calculated. To control for daily variation in incorporation rate, the values obtained on each day were normalized to the 35S-Met/Cys ACSF used for incubation, and the average incorporation of all slices analyzed in that experiment, as described^2^.

**Hippocampal slice electrophysiology.** Horizontal hippocampal slices (400 μm) were prepared from P25–32 male WT and *FMR1^CRE^* knock-in mutant litter mate mice. Slices were collected in carbogenated (95% oxygen, 5% CO2) ice-cold dissection buffer containing the following (in mM): 86 NaCl, 1.2 NaH2PO4, 25 KCl, 25 NaHCO3, 20 glucose, 75 sucrose, 0.5 CaCl2, and 7 MgCl2. Slices were incubated for 30 min at ∼30°C in artificial CSF (ACSF) containing the following (in mM): 124 NaCl, 1.2 NaH2PO4, 25 KCl, 25 NaHCO3, 20 glucose, 2 CaCl2, and 1 MgCl2, bubbled with 95% oxygen and 5% CO2. An incision was made through CA1–CA3 boundary, and slices were left to recover for a minimum of 1 h at room temperature (20−22°C) before any recordings were made. For electrophysiological recordings, slices were placed in a submersion chamber heated to 30°C and perfused with carbogenated ACSF at a rate of 4 ml/min.

Field EPSPs (fEPSPs) were recorded at Schaffer collateral/commissural inputs to CA1 pyramidal neurons using extracellular recording electrodes (1–3 MΩ) filled with ACSF and placed in the stratum radiatum layer of the CA1 area. Synaptic responses were evoked by applying single current pulses to the Schaffer collateral/commissural axons using a bipolar stimulating electrode. Stimuli (200μs duration, 30s interval) were set to produce 40–60% of the maximal response amplitude. Metabotropic glutamate receptor-dependent LTD (mGluR-LTD) was induced by acute application (5 min) of the group 1 mGluR agonist 3,5-dihydroxyphenylglycine (DHPG; 100 μM).

Electrophysiological traces were collected using WinLTP (University of Bristol, Bristol, UK) and exported to Microsoft Excel. The magnitude of LTD was calculated by dividing the average fEPSP slope from 50 to 60 min after DHPG application by the average fEPSP slope during the 20 min baseline before DHPG application. Statistical analysis was performed using GraphPad Prism (GraphPad Software). Time-matched normalized data were averaged across experiments and expressed as means ± SE. Significant differences between the WT and Fmr1CRE knock-in mutant mice were determined using Student's t test.

**RNA isolation and quantitative PCR.** Total RNA was extracted from dissected fore brain, mid brain and hind brain of 3-4 biological replicates of P-7,P-14 and P-25 wild type and *Fmr1*^CRE^ knockin mutant mice from same litter using the RNeasy kit (QIAGEN) combined with QIAshredder (QIAGEN), following the manufacturer’s instructions. cDNA was synthesized from 1 μg of total RNA using the Roche First Strand Synthesis Kit. Quantitative PCR was performed using the Roche LightCycler 480 and the Roche LightCycler 480 Probes Master. All samples were analysed in biological triplicates or quadruplets and technical triplicates. Primer sequences are described in **Table S2** and probe numbers for *Fmr1* (6 and 78) and *Gapdh* (52) were used .*Fmr1* transcript levels were normalized to *Gapdh* levels for all experiments.

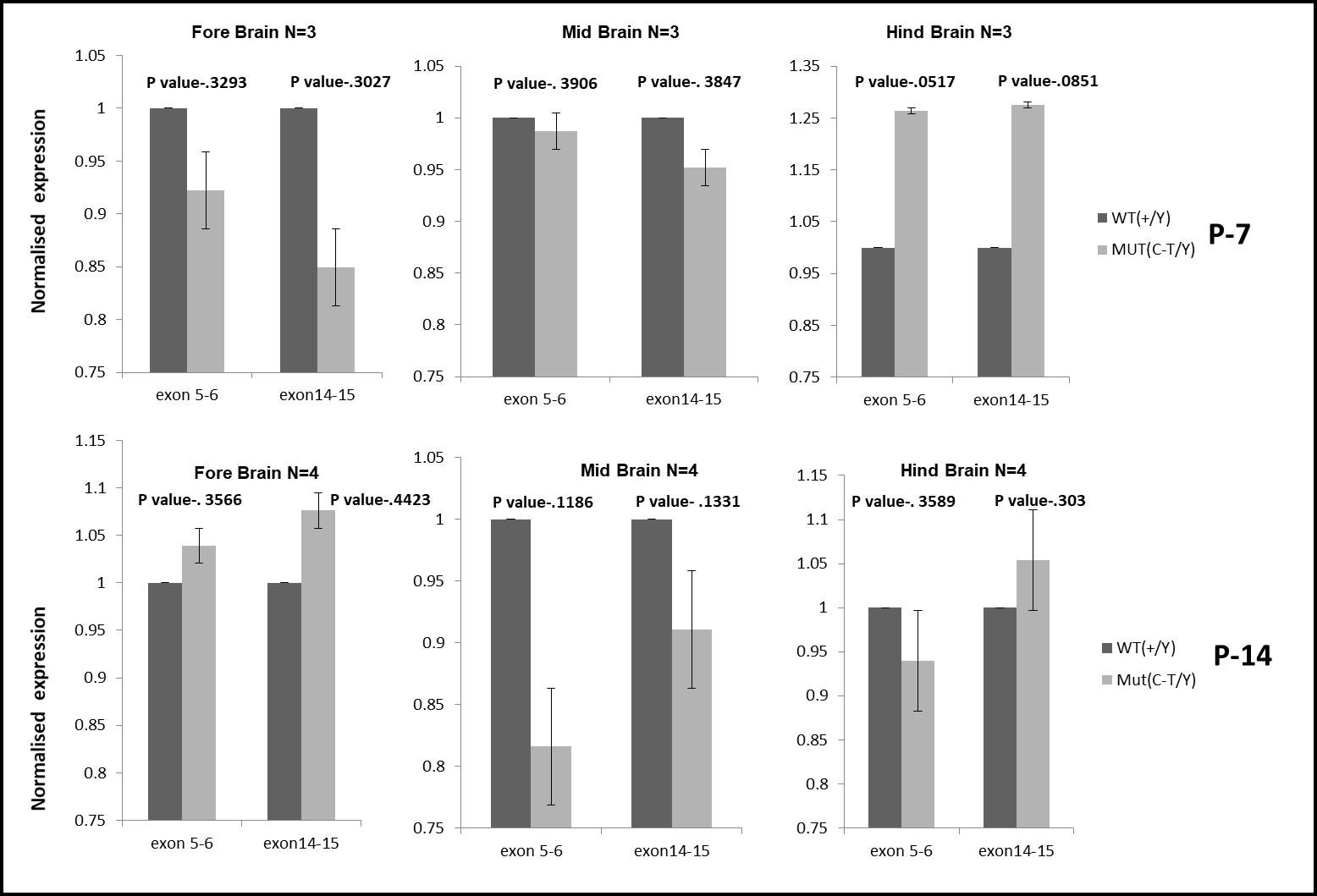

**Supplementary Fig. 2:** **Quantification of *Fmr1* transcripts using quantitative PCR.** Transcripts level of *Fmr1* is quantified and normalised with *Gapdh* transcripts levels in the forebrain, midbrain and hindbrain of *Fmr1*^CRE^ knockin mutant as compared to wild type littermates at different stage of development (P7 and P14).Two different exonic regions were used to compare the *Fmr1* transcripts levels. The levels of *Fmr1* transcripts were not significantly different between *Fmr1*^CRE^ knockin mutant as compared to wild type littermates in different tissues as well as postnatal developing stages (P-7 and P-14)

**RNAscope assay.** Whole brain were dissected out from male WT and *Fmr1*^CRE^ knock-in mutant mice(P-25) followed by fixation with 4% paraformaldehyde and processed for cryostat sections (20 μm). The Sagittal sections were subjected to RNAscope Multiplex Fluorescent Assay. The sections were incubated with pretreatment 2 (RNAscope Target Retrieval Reagents, Advanced Cell Diagnostics) for 30 minutes at 40°C in a HybEZ oven (HybEZ Hybridization System, Advanced Cell Diagnostics), followed by pretreatment 3 (RNAscope Protease III, Advanced Cell Diagnostics) containing protease for 30 minutes at 40°C. After pretreatment 3, the sections were washed with 5× deionized water and incubated with the prewarmed mixed mRNA target probes (FMR1- C2, 496399 and PAX-6-C1, 412821 [Advanced Cell Diagnostics]) for 2 hours at 40°C in the HybEZ oven. The sections were processed in wash buffer for 5 minutes and then incubated with Amp1 for 30 minutes at 40°C and washed with washing buffer for 5 minutes. The procedure was repeated for Amp2, Amp3, and Amp4 AltA for C2 (Cy5), and C1 (FITC) module, After washing with washing buffer for 5 minutes, the sections were cover slipped with Vectashield-DAPI mounting medium (Vector Laboratories).

**RNA Sequencing.** Total RNA was extracted from dissected fore brain, mid brain and hind brain of 3 biological replicates P-25 wild type and *Fmr1*^CRE^ knockin mutant mice from same litter using the RNeasy kit (QIAGEN) combined with QIAshredder (QIAGEN), following the manufacturer’s instructions.For RNA-sequencing, random primed cDNA from poly(A) selected RNA was converted into an Illumina sequencing library using RNA Library Prep Kit from Illumina (E7420, NEB, USA) in conjunction with NEBNext® Multiplex Oligos for Illumina (E7335/E7500, NEB, USA). and single-end 50-base pair (bp) reads were generated using a NextSeq 500 (Illumina Inc, SY-415-1002). Eighteen libraries were combined in two equimolar pools of 9 based on the library quantification results and each pool was run across a single High-Output Flow Cell. Sequencing was performed at the Wellcome Trust Clinical Research Facility (WTCRF; Edinburgh).

Fastq files were processed to transcript-level counts and quality control performed using the bcbio_nextgen pipeline and the illumina-RNAseq workflow template. Differential Expression (DE) analysis was performed in R. The R package bcbio-RNAseq was used to import salmon transcript level counts into DESeq2.

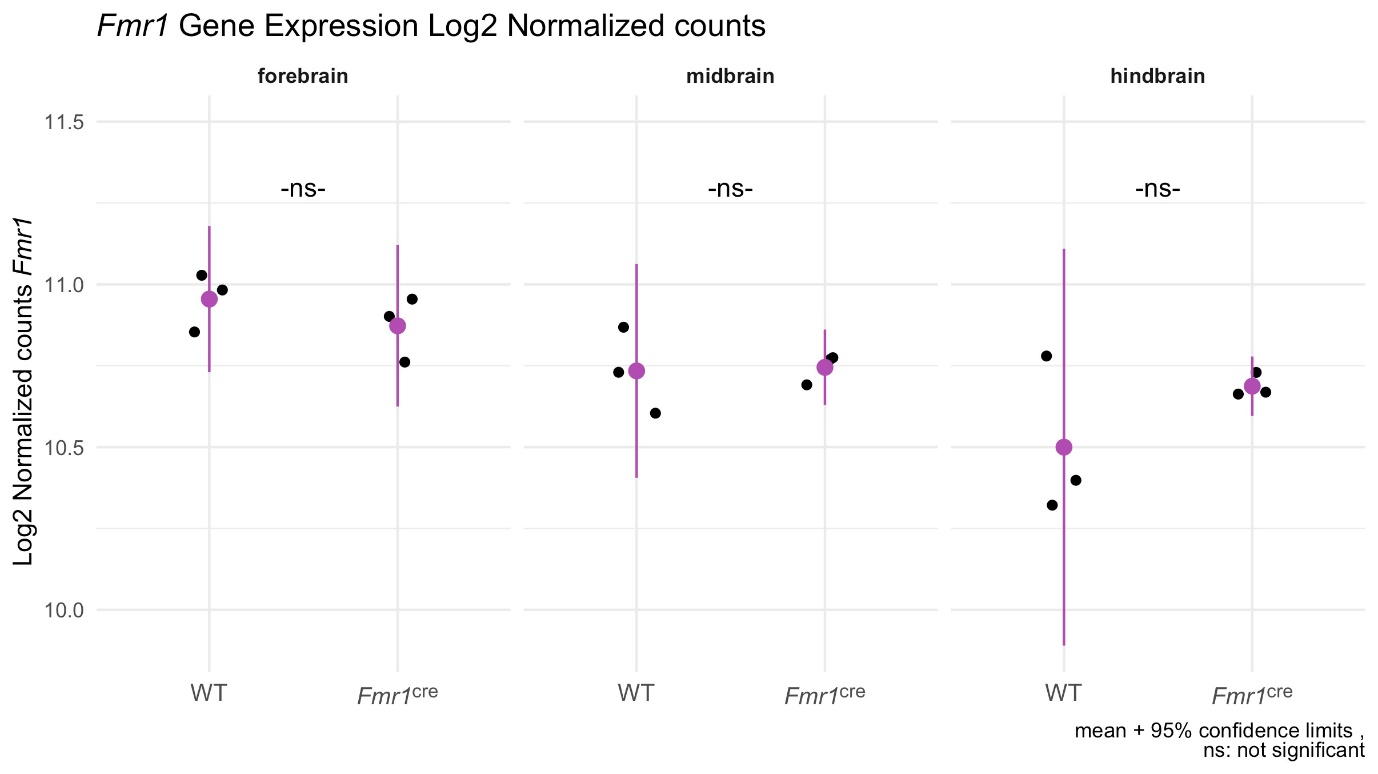

**Supplementary Fig. 3:** ***Fmr1* transcripts counts using RNA Sequening.** DESeq2 was used to quantify the Log2 normalised *Fmr1* transcripts levels in the forebrain, midbrain and hindbrain of 3 biological replicates of *Fmr1*^CRE^ knockin mutant as compared to wild type littermates The Log2 normalised *Fmr1* transcripts counts were not significantly different between *Fmr1*^CRE^ knockin mutant as compared to wild type littermates in forebrain, midbrain and hindbrain at P-25 stage of development.

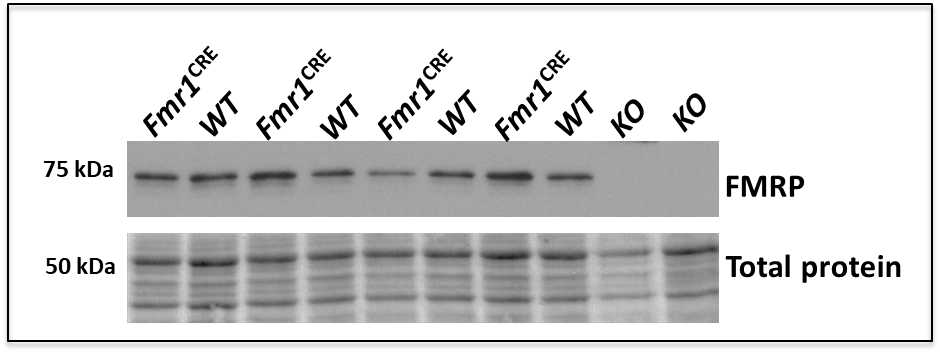

**Supplementary Fig. 4: FMRP expression in the hippocampal slices.** Western blotting was performed to detect the levels FMRP in the hippocampal slices of *Fmr1*^CRE^ knockin mutant as compared to wild type litter mates. Samples from KO (*Fmr1* knockout) was used as negative control samples. Total protein in a comassie stain gel was used as loading control.The levels of FMRP was quantified and normalised against total proteins. (Fig. 4b)

**Supplementary Note:**

**Details of Guide RNA and Repair template Sequence used in the study**

**gRNA *Tenm1*^CRE^***:* AATATTATTAGCCACACATT **TGG**

**Repair template *Tenm1*^CRE^:** AAAGATTGAAATAATTTTATAGAAAGCTTAAAAGTTAAATTCTTCATTATAAATCTCATTGGTTATATGTAGAGTTTCAGATAAACAGAATGACTGTCGAATGATATTTACCAAATGTGTGGCTAATAATATTCCCCCACTTCTAAATATGCATCACTTTTATTGACATAAAAAATATAT

**gRNA *Fmr1*^CRE^***:* ACCTTGTGTCTATGACTATT **TGG**

**Repair template *Fmr1*^CRE^:** TAAGATGGATTCATATTAGGGCTCAAATGCATTGATAGCATTCTACATATTTTTATCCATTTTTATTCCAAGCTACTTTTATCCAAATAGTTATAGACACAAGGTTATTGCAAATTGTATTTGTCTGCTGCCATAGTGCTTTCTATTTTAGAGGAGTAGAAGTAACTATCTCCTTAACAA

Guide RNA and Repair template sequence used to create human variant in *Tenm1* and *Fmr1* CRE in mouse. PAM sequence in the guide RNA is marked in bold and variant nucleotide in repair template is highlighted.
