## Supplementary Fig for "Functional Predictors of Causative *Cis*-Regulatory Mutations in Mendelian Disease"

Family 391, chrX:g.147866225C>A, the variant was confirmed in the studied individual; DNA from other family members was not available for segregation

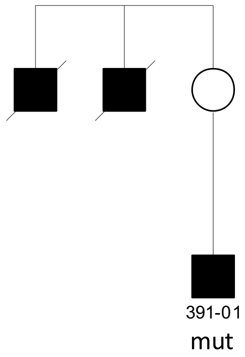

Family 061, chrX:g.99625573T>A, the variant did not segregate as expected in the family

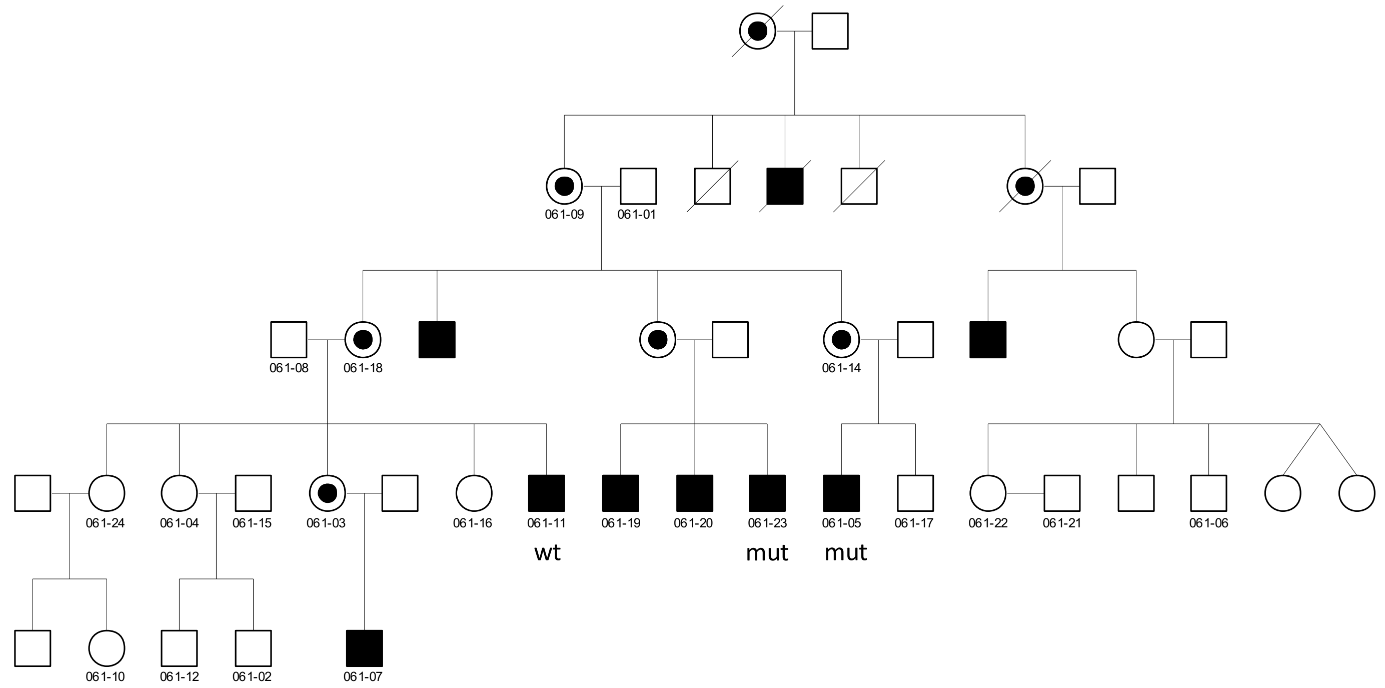

Family 136, chrX:g.136354272G>A, the variant did not segregate as expected in the family

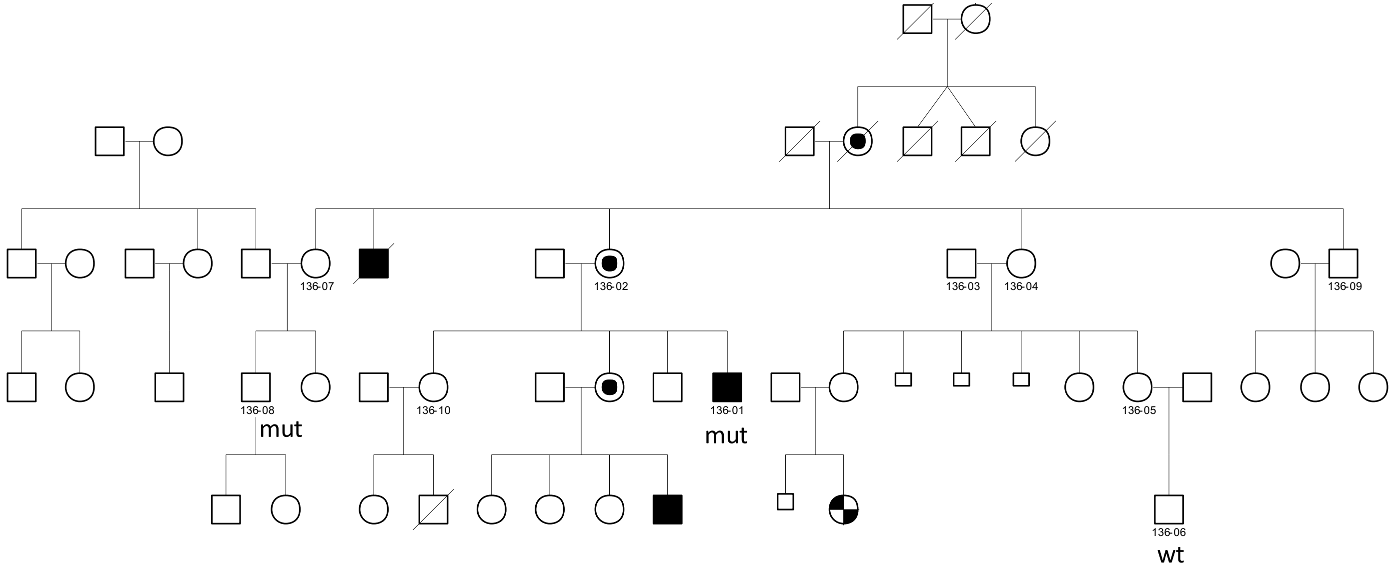

Family 061,chrX:g. 135962942G>C, the variant did not segregate as expected in the family

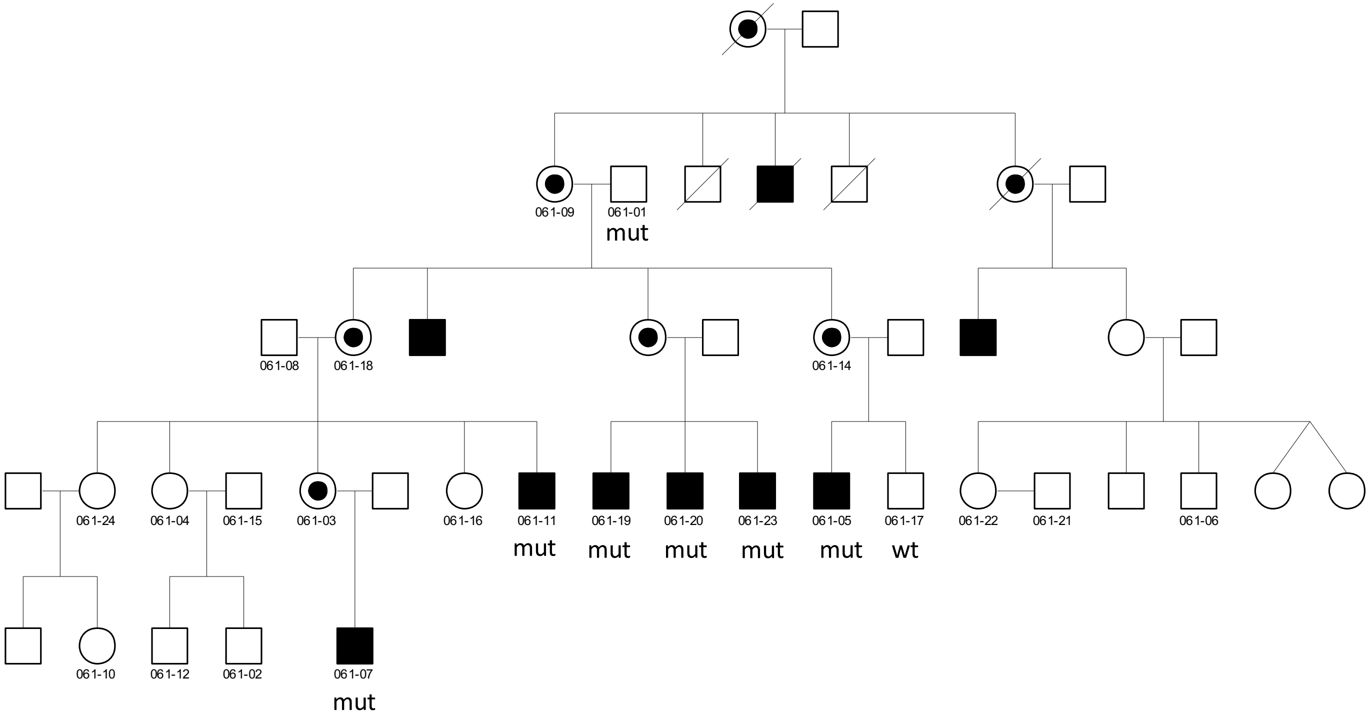

Family 375, chrX:g. 24819218G>C, the variant did not segregate as expected in the family

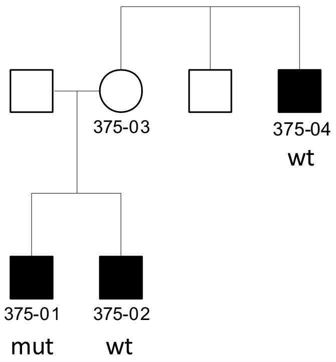

Family 102, chrX:g.148000915A>G, the variant did not segregate as expected in the family

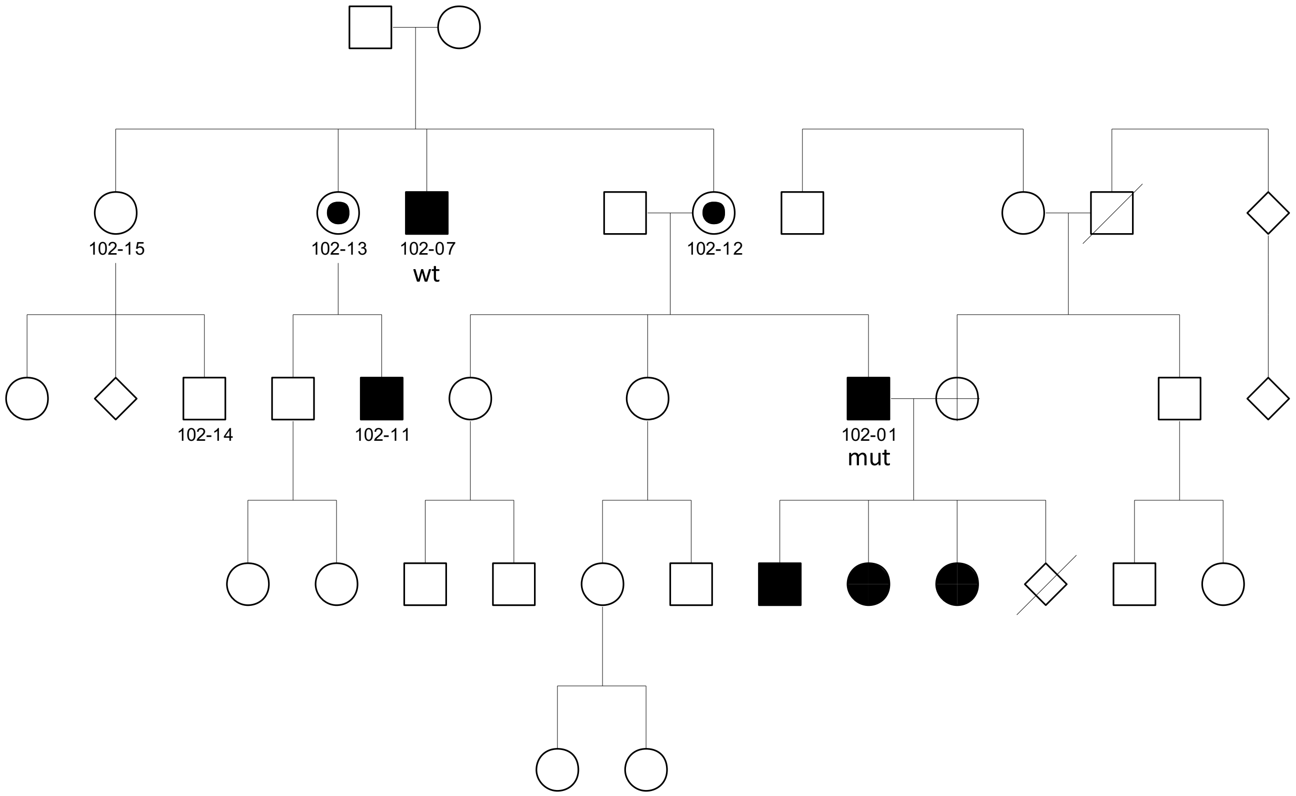

Family 409, chrX:g.136183176G>A, the variant segregated as expected in the family

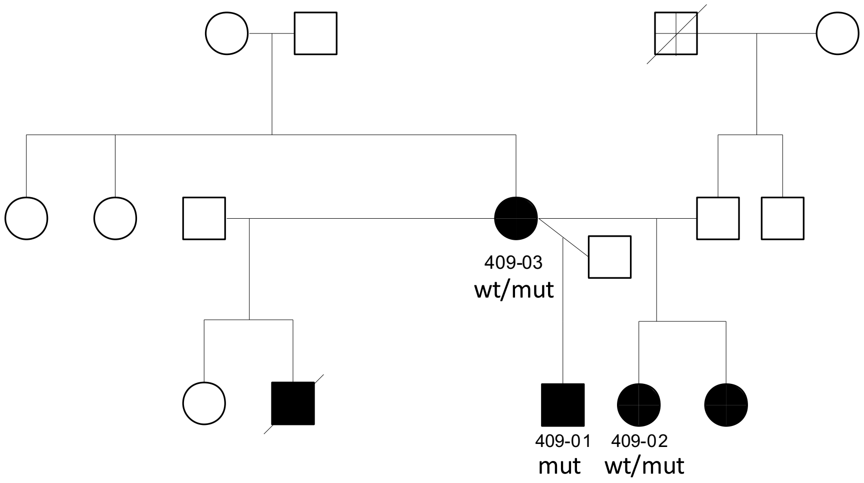

Family 401, chrX:g.123707760T>C, the variant did not segregate as expected in the family

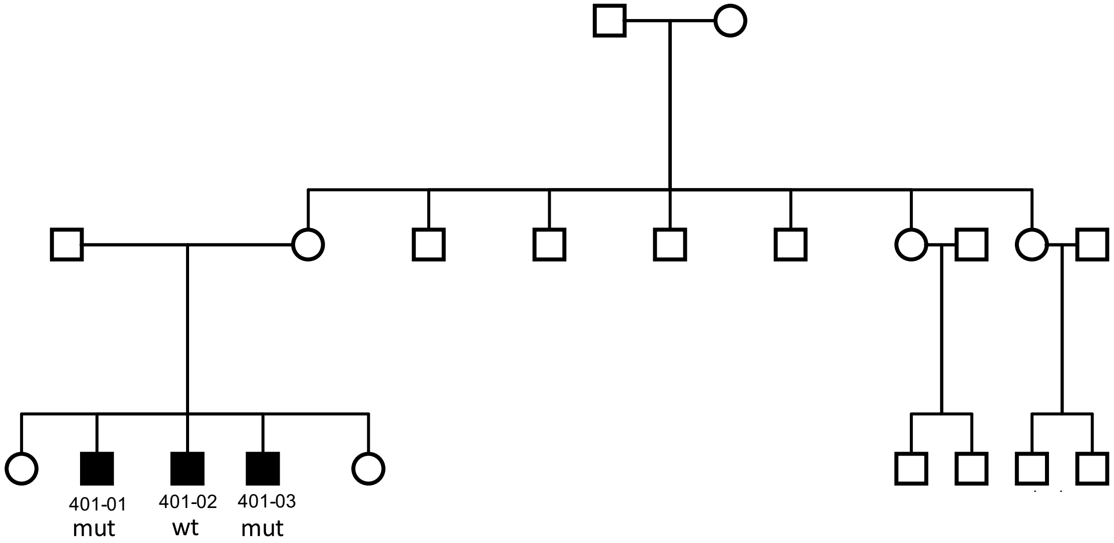

Family 006, chrX:g.139187167A>G, the variant did not segregate as expected in the family

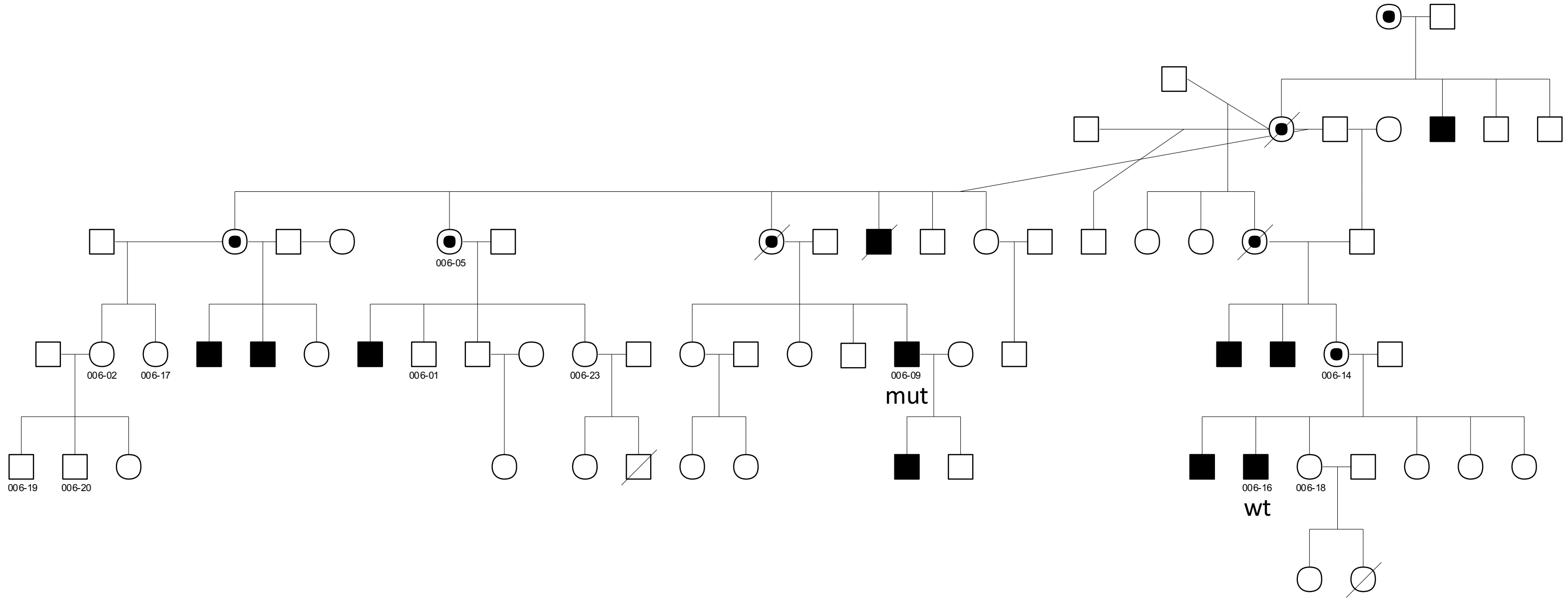

Family 083, chrX:g.39701238C>G, the variant did not segregate as expected in the family

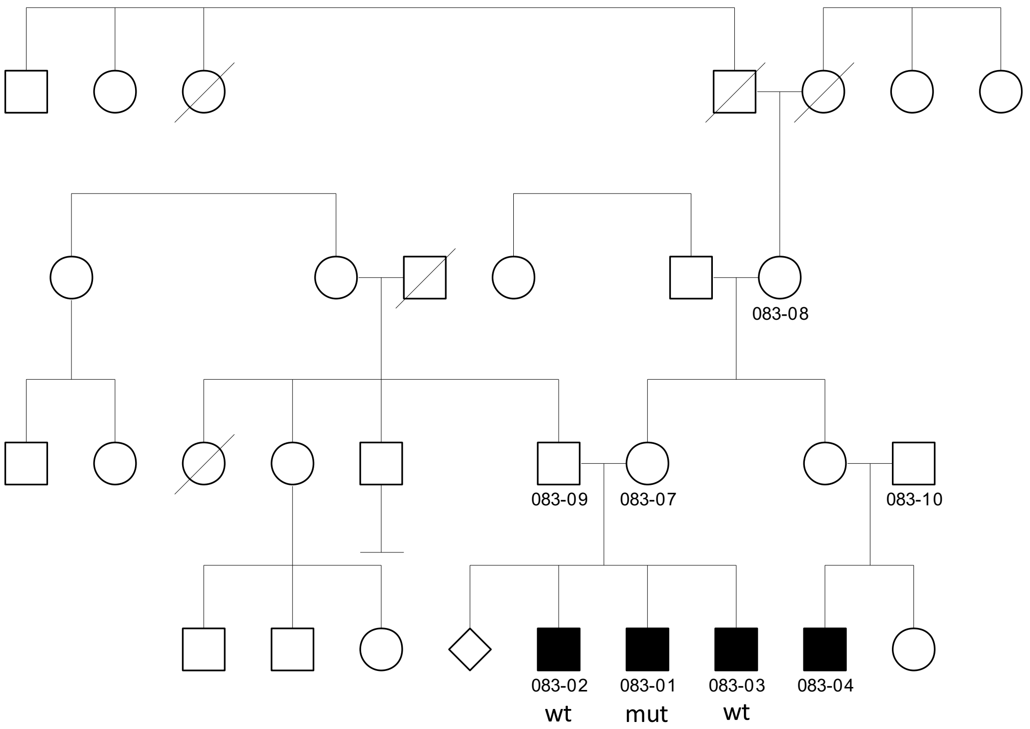

Family 375, chrX:g.40406651T>C, the variant did not segregate as expected in the family

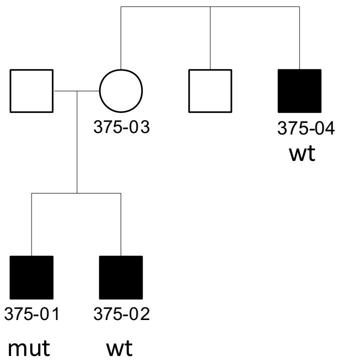

Family 136, chrX:g.25260740A>G, the variant segregated as expected in the family

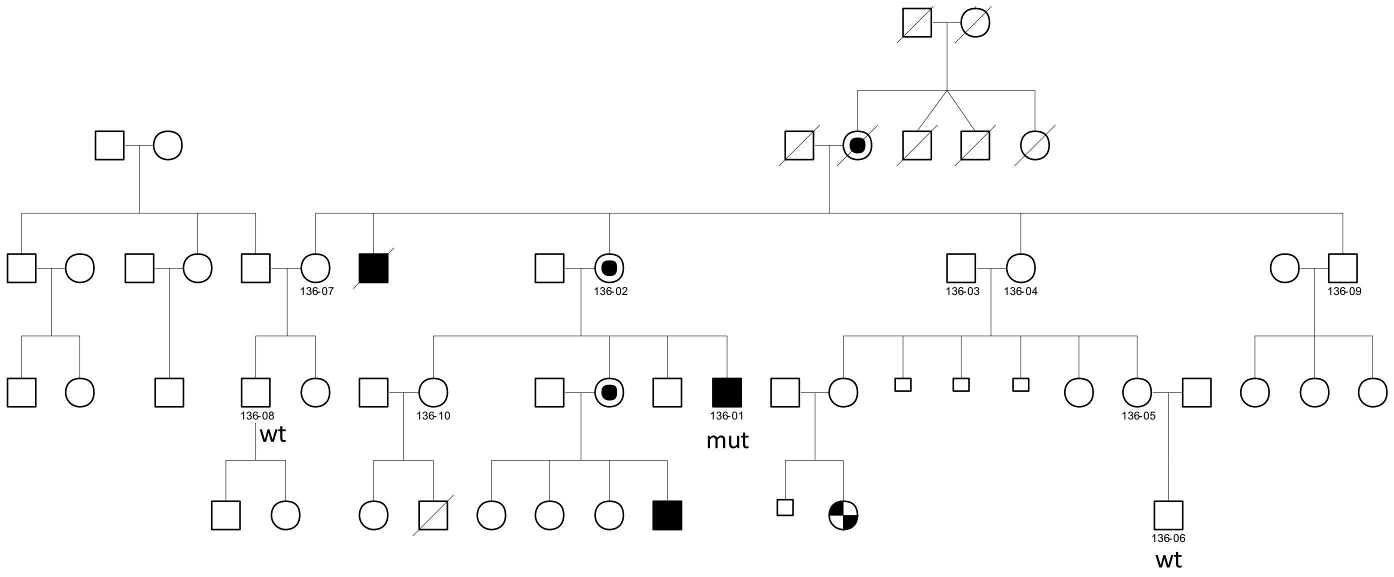

Family 029, chrX:g.106176958A>C, the variant was not confirmed with Sanger sequencing

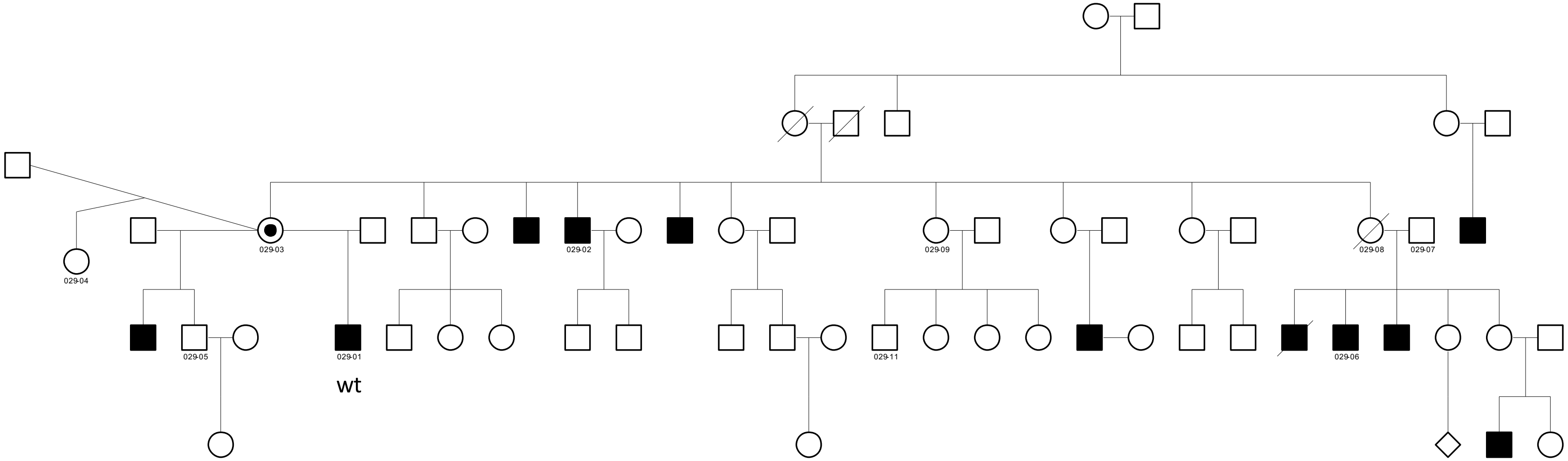

Family 083, chrX:g.85932752A>G, the variant did not segregate as expected in the family

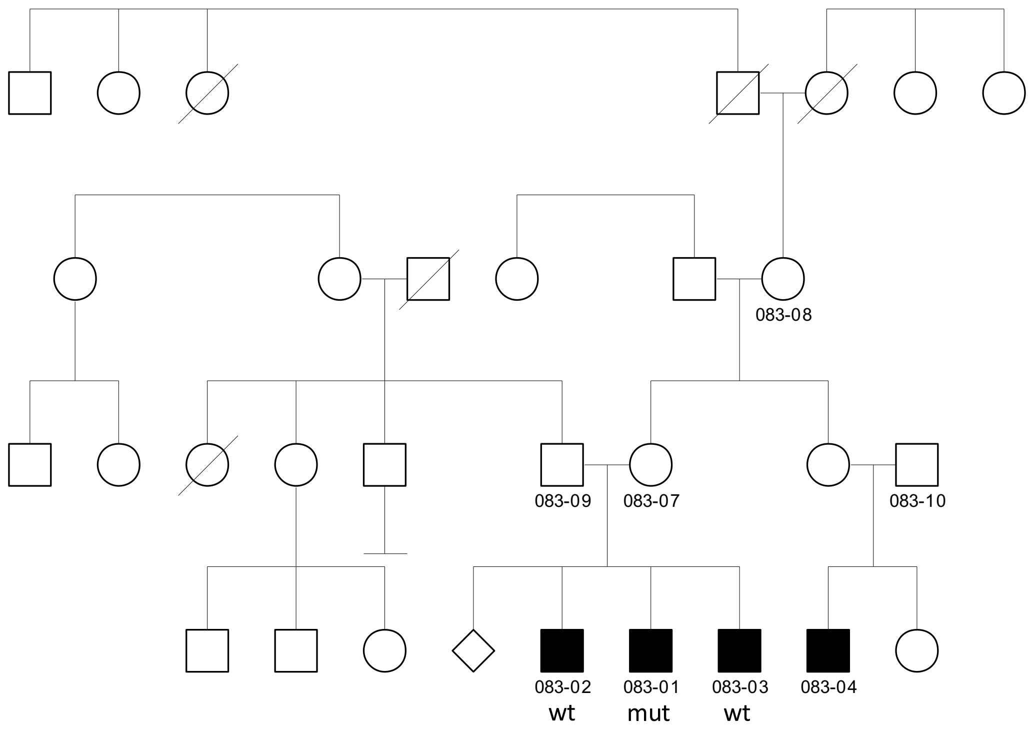

Family 375, chrX:g.33131770G>C, the variant did not segregate as expected in the family

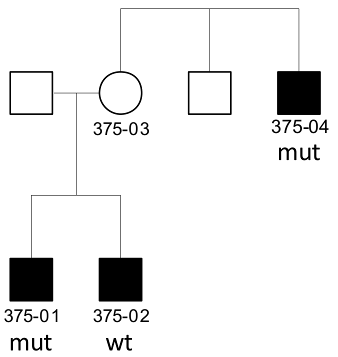

Family 409, chrX:g.124269322G>A, the variant segregated as expected in the family

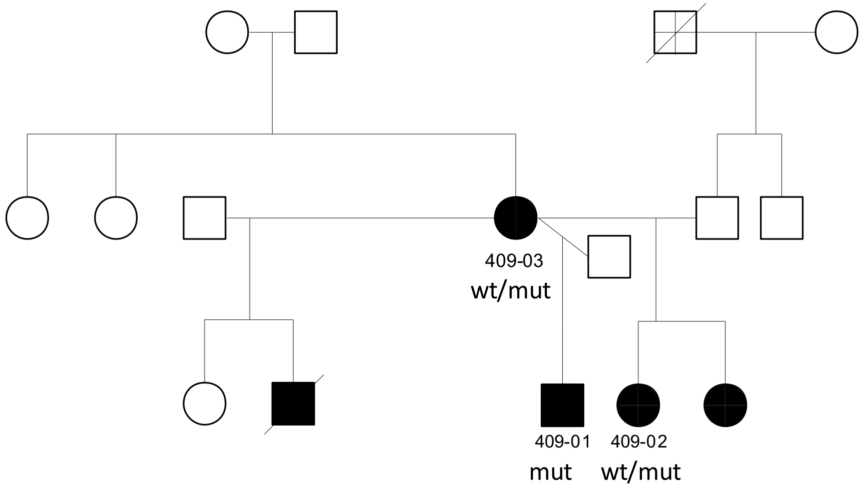

Family 006, chrX:g.15010623G>A, the variant did not segregate as expected in the family

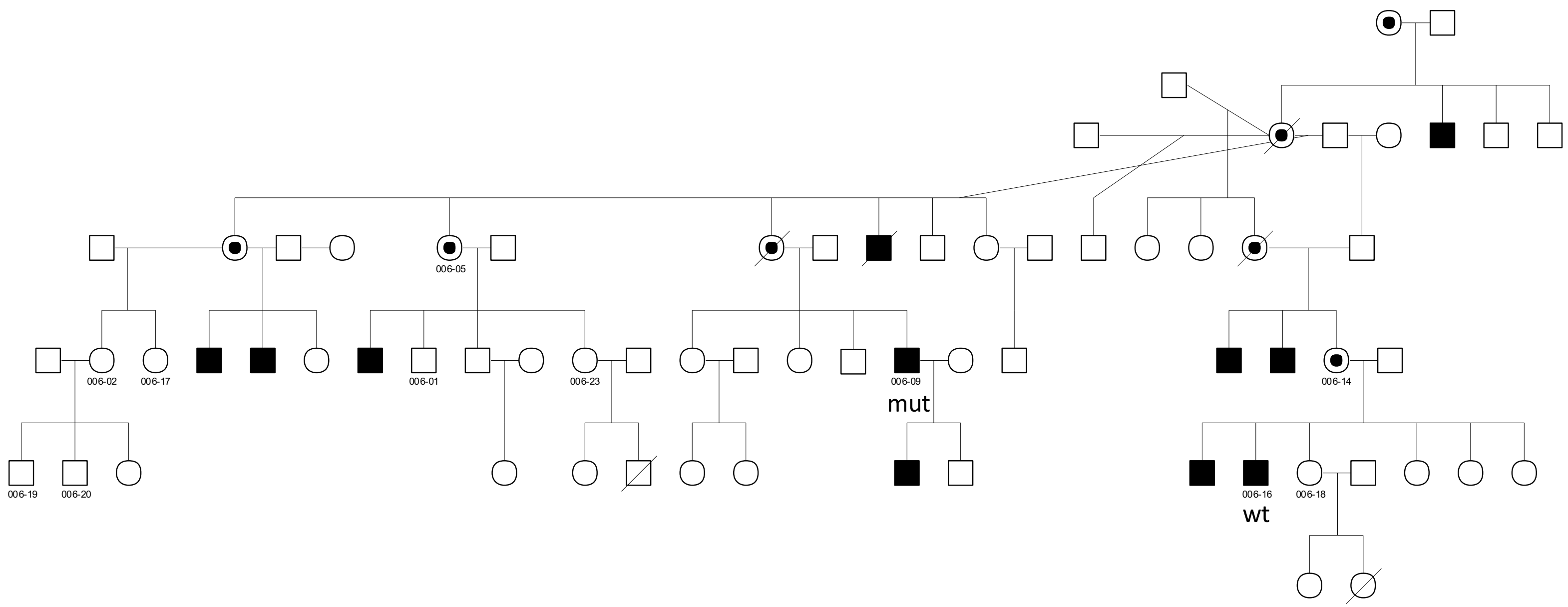

Family 351, chrX:g.135962942T>C, the variant did not segregate as expected in the family

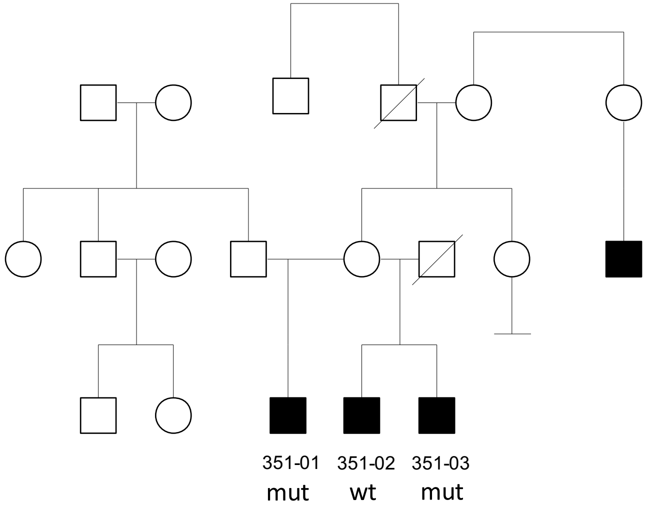

Family 391, chrX:g.104365258A>G, the variant was confirmed in the studied individual; DNA from other family members was not available for segregation

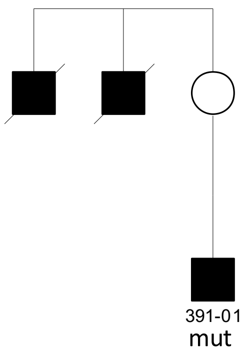

Family 010, chrX:g.116741775C>T, the variant did not segregate as expected in the family

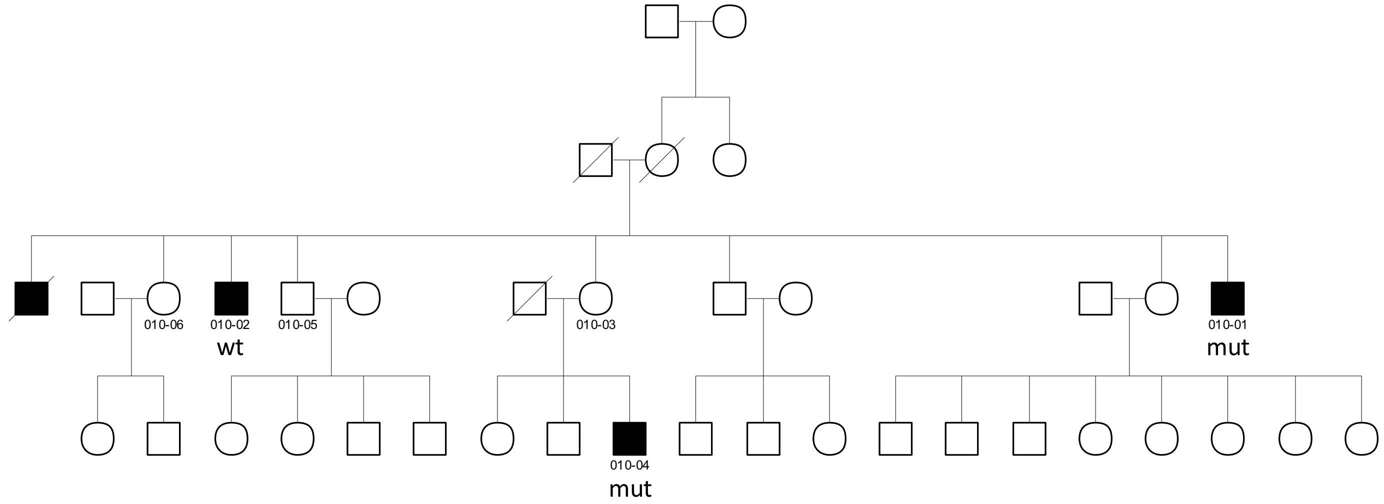

Family 347, chrX:g.146875009C>T, the variant segregated as expected in the family

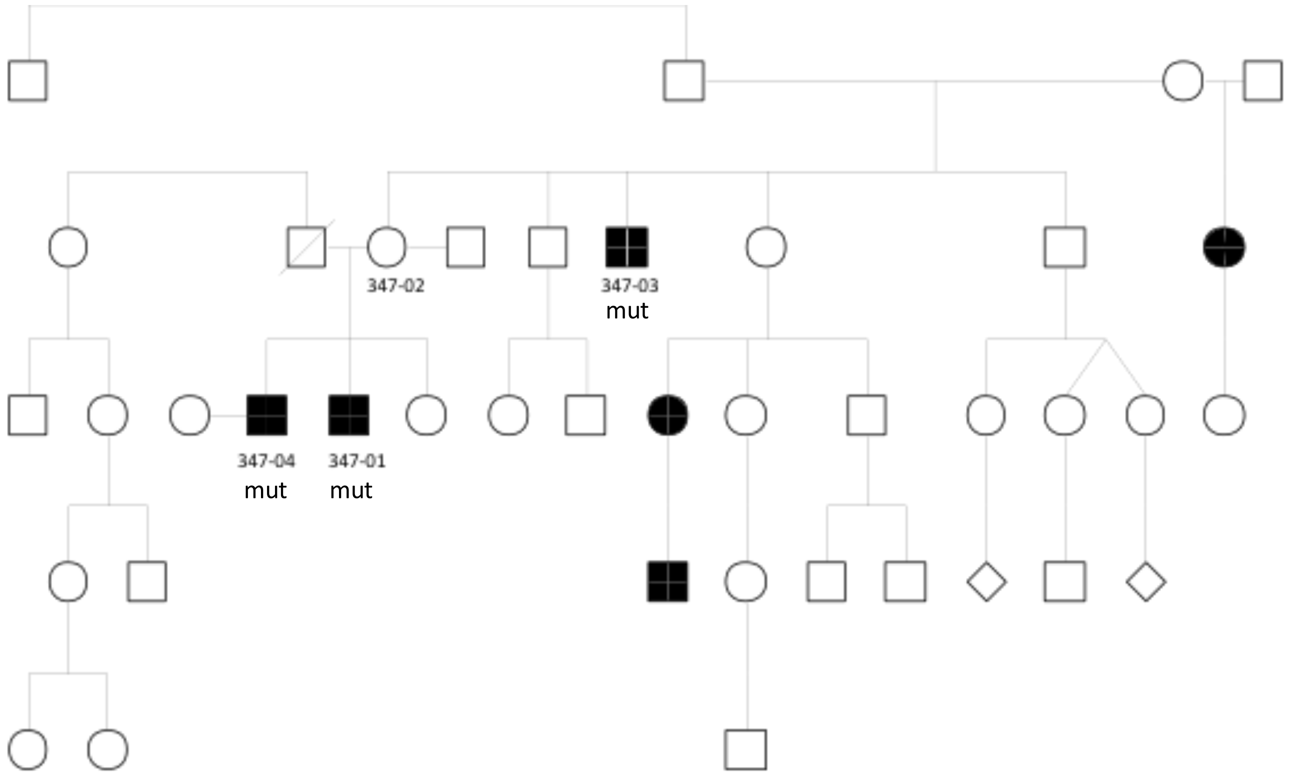

Family 440, chrX:g.103994885G>A, the variant was confirmed in the studied individual; DNA from other family members was not available for segregation

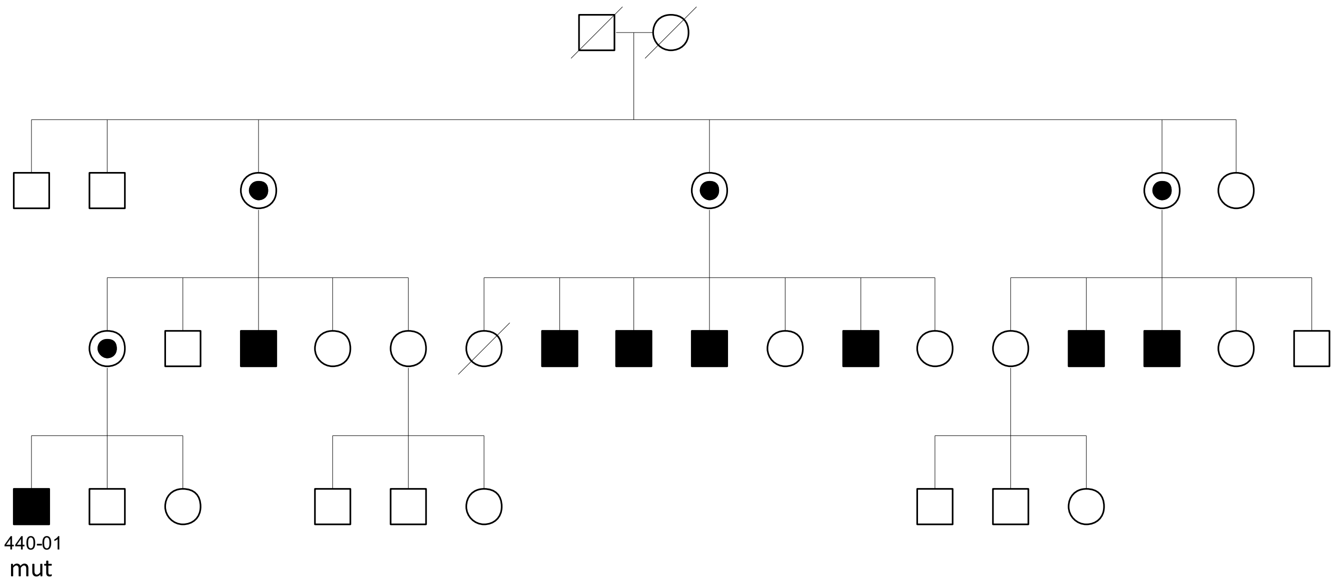

Family 370, chrX:g.133105579G>A, the variant did not segregate as expected in the family

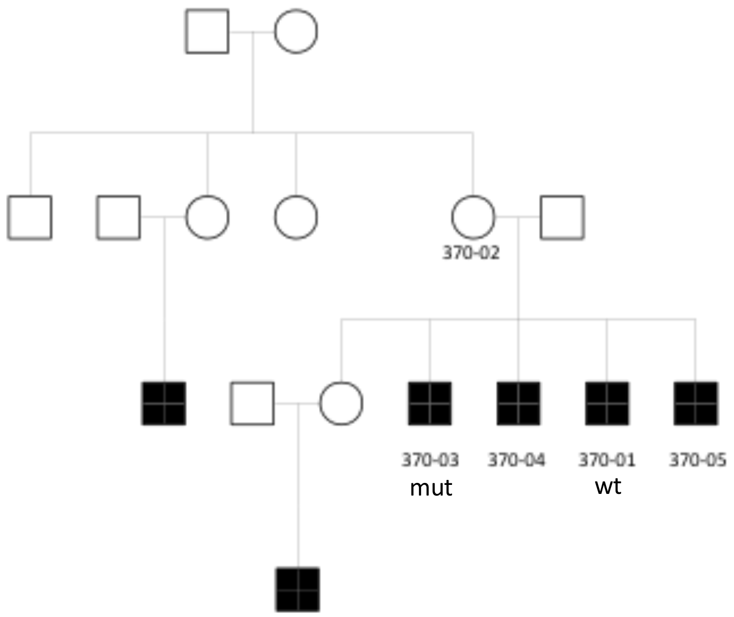

Family 383, chrX:g.45375111C>G, the variant segregated as expected in the family

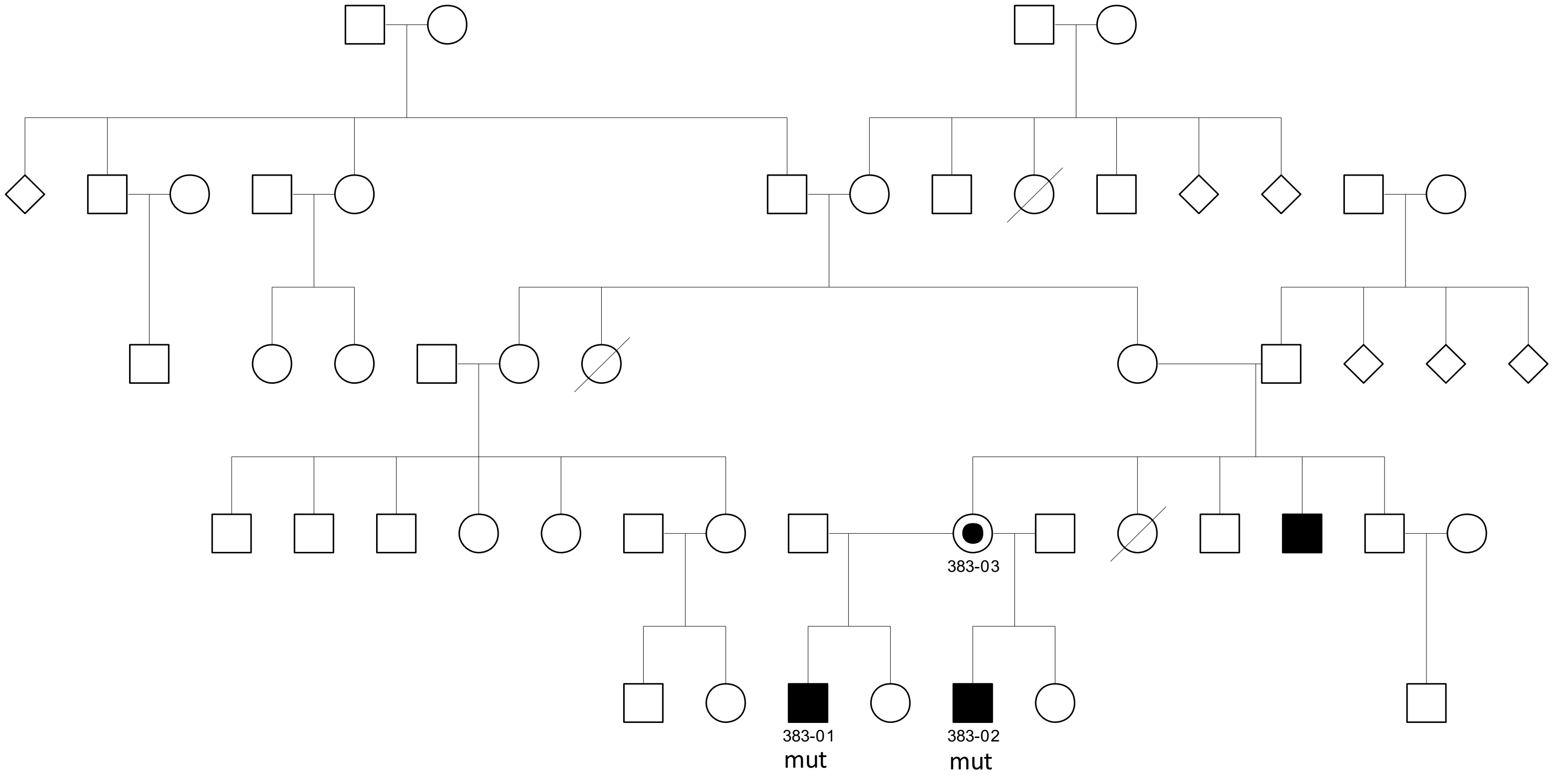

Family 382, chrX:g.135112643T>G, the variant did not segregate as expected in the family

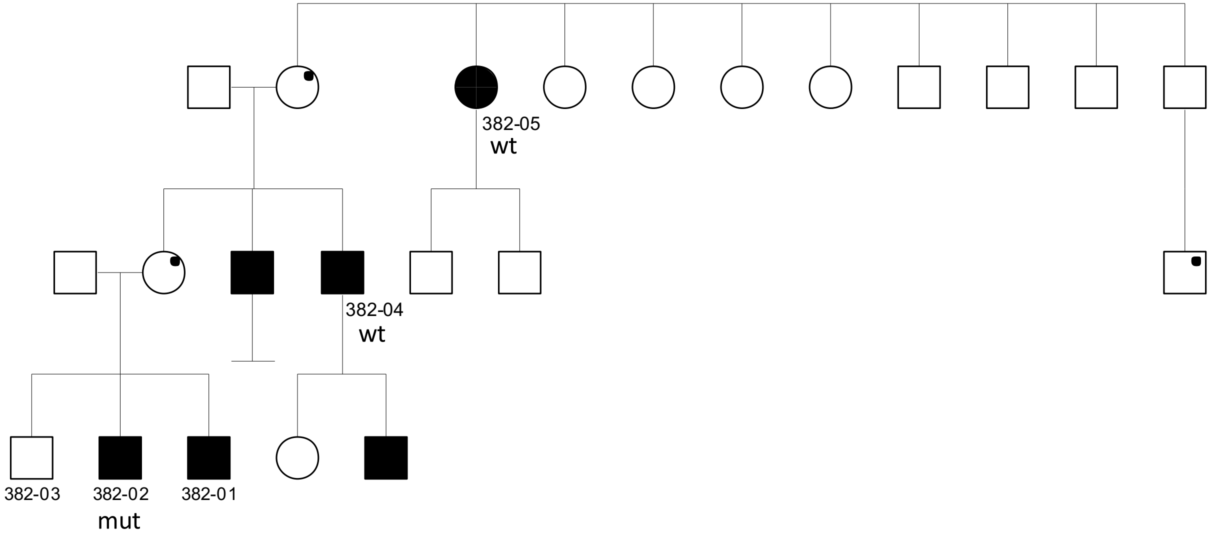

Family 1319, chrX:g.39719711T>C, the variant was confirmed in the studied individual; DNA from other family members was not available for segregation

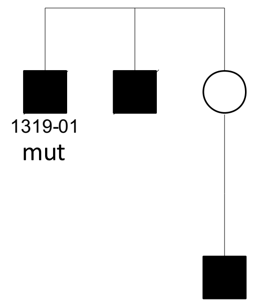

Family 439, chrX:g.38741210T>A, the variant was confirmed in the studied individual; DNA from other family members was not available for segregation

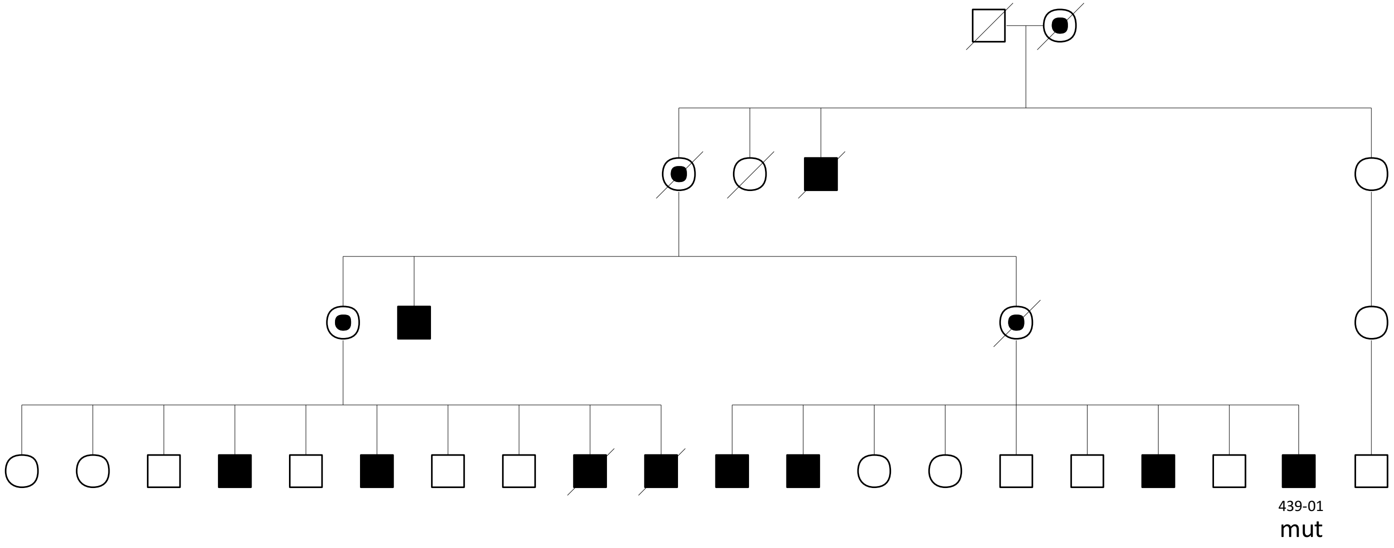

Family 429, chrX:g.39976350C>G , the variant did not segregate as expected in the family

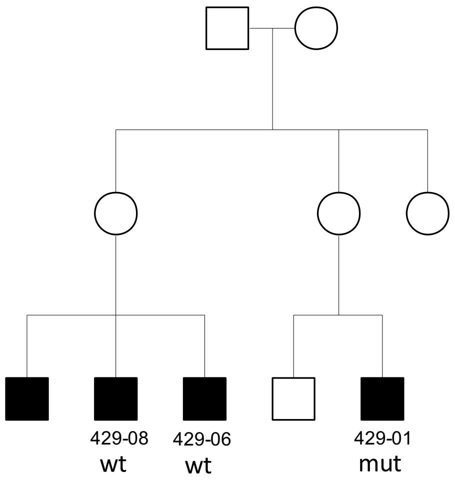

Family 438, chrX:g.128915530C>G, the variant was confirmed in the studied individual; DNA from other family members was not available for segregation

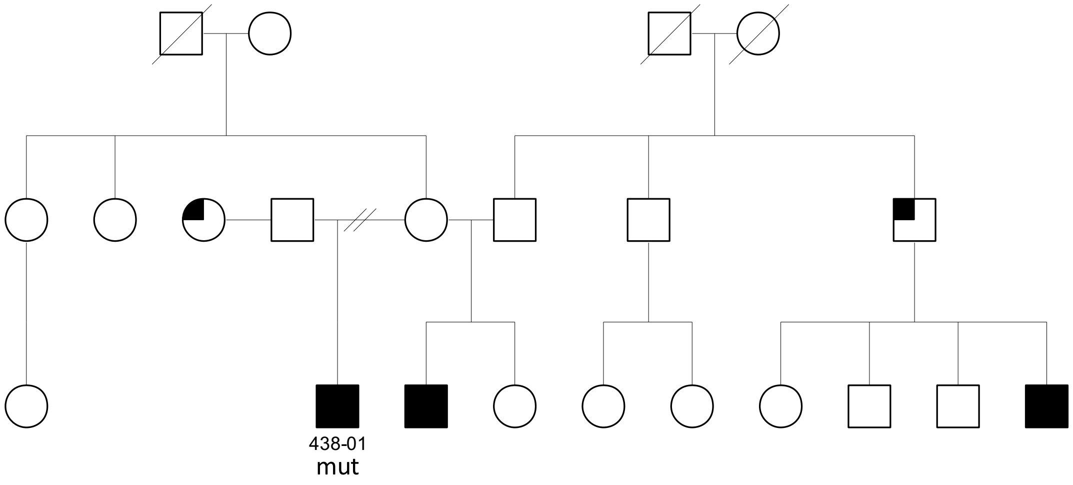

Family 083, chrX:g.39377898A>G, the variant did not segregate as expected in the family

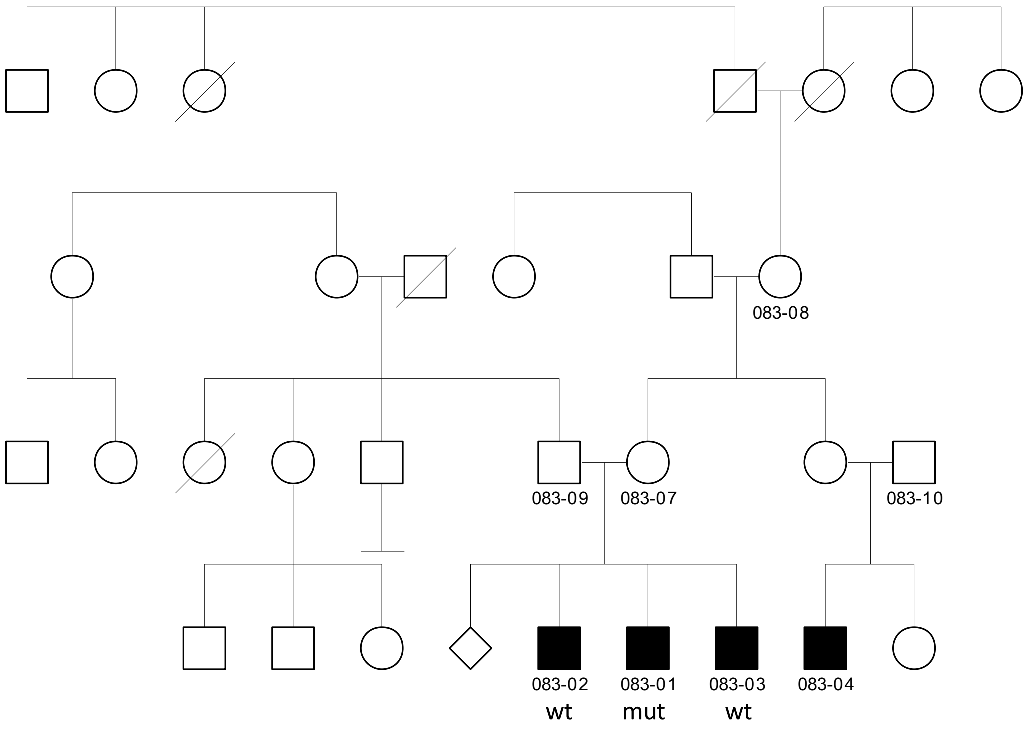

Family 375, chrX:g.34895739T>C, the variant did not segregate as expected in the family

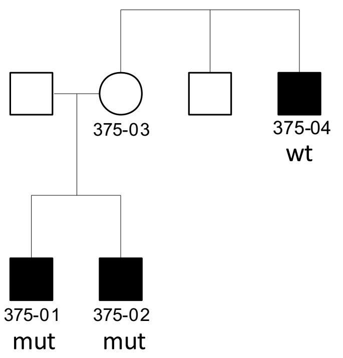
